# Regeneration of humoral immunity from pluripotent stem cells by defined transcription factors

**DOI:** 10.1101/2021.07.30.454442

**Authors:** Qi Zhang, Bingyan Wu, Qitong Weng, Fangxiao Hu, Yunqing Lin, Chengxiang Xia, Huan Peng, Yao Wang, Xiaofei Liu, Lijuan Liu, Jiapin Xiong, Yang Geng, Yalan Zhao, Mengyun Zhang, Juan Du, Jinyong Wang

## Abstract

Regeneration of humoral immunity from pluripotent stem cells (PSCs) is a crucial aim in translational medicine. However, reconstitution of complete, sustained, and functional B lymphopoiesis from PSCs has not yet been developed. Here, we successfully achieved regenerative B lymphopoiesis in B-cell deficient animals transplanted with PSC-derived hematopoietic progenitors (iHPCs) guided by synergistic expression of *Runx1, Hoxa9*, and *Lhx2*. Upon transplantation, the iHPCs immediately gave rise to pro/pre-B cells in recipients’ bone marrow, which were able to further differentiate into the entire B cell lineages, including innate B-1a, B-1b, MZ B cells, as well as adaptive FO B cells. In responding to antigen stimuli, the regenerative B cells produced adaptive humoral immune responses, sustained a prolonged antigen-specific antibody production, and formed immune-memory. Particularly, the regenerative B cells in spleen showed developing patterns of immunoglobulin chain-switch and hyper-mutation via a cross-talk with the host T follicular helper cells, which eventually formed T cell-dependent humoral responses. This study provides *de novo* evidence that B lymphopoiesis can be regenerated from PSCs via a HSC-independent approach, which provides insights into treating B-cell related humoral deficiencies using PSCs as unlimited cell resource.

## INTRODUCTION

B cells, including invariant B-1 cells and adaptive B-2 cells, are the essential cell types of humoral immune system. B-1 cells maintain self-renewal capacity in peritoneal and pleural cavities of adults and perform a rapid innate immune response against common pathogens by producing natural antibodies (Baumgarth, 2011). Adult HSCT poorly reconstitutes B-1a cells (Barber et al., 2011; Ghosn et al., 2012), indicating an adult HSC-independent origin. B-2 cells, including follicular B (FO B) cells and marginal zone B (MZ B) cells, mature in spleen and are essential for adaptive humoral responses. FO B cells are in charge of specific adaptive immune response against foreign antigens by producing high-affinity antibodies and generating immunological memory in a T cell-dependent manner (Allman and Pillai, 2008), MZ B cells are innate-like antibody-producing cells which locate at the interface between the circulation and the white pulp of the spleen, where provides a first line of defense against blood-borne viruses and encapsulated bacteria (Cerutti et al., 2013; Martin et al., 2001). Absence or reduction of any subset of B cells can lead to a compromised humoral response and severe infections from bacteria, viruses, and other microbes (Smith and Baumgarth, 2019; Vale and Schroeder, 2010; Zouali and Richard, 2011). To overcome these B-cell related defects, regeneration of B cells is an ideal option.

Bone marrow microenvironment accounts for supporting proliferation, differentiation, and survival of early B cell precursors with typical molecular events of immunoglobulin heavy chain VDJ recombination, pre-BCR, and BCR checkpoints (Melchers, 2015). Surface IgM^+^ (sIgM^+^) immature B cells leave the bone marrow and settle in the spleen, wherein B cell mature and survive via complicated and kinetic positive signals (Mackay and Schneider, 2009). In the presence of T cell-dependent antigen, the spleen germinal center (GC) is established and polarized into two compartments known as the dark zone and the light zone. The network of CXCL12-producing reticular cells (CRCs) in dark zone provides the structural and chemotactic support for activated B cells localization and proliferation and promotes the process of somatic hypermutation. The tight interactions between activated B cells and multiple cell types in light zone, particularly follicular dendritic cells (FDC) and T follicular helper cells (Tfh), drive the processes of high affinity maturation and class switch to produce high affinity antibodies (De Silva and Klein, 2015; Stebegg et al., 2018; Vinuesa et al., 2010). Thus, consecutive B cell differentiation, maturation, and activation require multiple spatiotemporal stages, which depends on strict and complex microenvironments.

Researchers attempted several approaches to produce B cells *in vitro*, despite the fact that there are no methods of mimicking the spatiotemporal microenvironments of B cell development. In the presence of MS5 stromal cells, CD34^+^ blood progenitors can differentiate into B cell precursors and IgM^+^ B cells *in vitro* (Hirose et al., 2001; Ohkawara et al., 1998). CD93^+^ B progenitor cells and functional IgM^+^ B lymphocytes can be generated *in vitro* from mouse ESC/OP9 co-culture system with the addition of exogenous Flt-3L (Cho et al., 1999). Human iPSCs co-cultured with stromal cells *in vitro* were able to differentiate into IgM^+^ B cells (French et al., 2015). Mouse ESC-derived pro/pre-B cells can transiently produce B-1 and conventional B cells in Rag deficient mice (Potocnik et al., 1997). More recently, a study demonstrated that the ESC-derived B progenitors gave rise to a long-term production of B-1b and MZ B cells while lacked the essential adaptive FO B cells (Lin et al., 2019). Similarly, incomplete B-cell populations were generated in recipients transplanted with ESC-derived c-Kit^+^ hematopoietic progenitors (Pearson et al., 2015). Of note, a conventional concept to regenerate engraftable B lymphopoiesis from PSC is to produce HSC-like cell intermediates with complete blood lineage potential (Lu et al., 2016; Sugimura et al., 2017). However, generating engraftable HSC-like cells *in vitro* is extremely inefficient (Stik and Graf, 2018). Nonetheless, an efficient approach of regenerating the entire subsets of functional B-1 and B-2 cells from PSCs has never been achieved.

Recent studies show that yolk sac (YS) and para-aortic splanchnopleura (P-Sp) cells can generate B-1 progenitors (Kobayashi et al., 2014; Yoshimoto et al., 2011). pre-HSC isolated from YS and P-Sp also are capable of producing B-1 and B-2 cells (Hadland et al., 2017; Kobayashi et al., 2019), signaling that the B-1 and B-2 cell fate are determined preceding the emergence of definitive HSC. Our group have recently reported that induced hemogenic endothelial progenitors (iHECs), derived from embryonic stem cells with inducible expression of *Runx1* and *Hoxa9*, can generate induced hematopoietic progenitor cells (iHPCs) preferentially contributing to functional T cells *in vivo* (Guo et al., 2020). Thus, regeneration of lymphopoiesis from PSCs can be achieved in the absence of regenerative HSCs.

In this study, we identify that synergistic expression of *Runx1, Hoxa9*, and *Lhx2* dominantly confers B cell lineage fate on PSC-derived iHPCs and leads to complete B lymphopoiesis *in vivo* following a differentiation scheme we previously reported (Guo et al., 2020; Wang et al., 2020). The regenerative B (iB) cells, including B-1a, B-1b, FO B, and MZ B subsets, possess a diversity of BCR repertoires similar to their natural B cell counterparts. These iB cells restored the B-cell deficient mice’s antibody responses boosted by specific antigen inoculation and maintained a long term humoral protection. For the first time, in the absence of iHSC, we establish a *de novo* approach of exclusively generating functional and complete B lymphopoiesis using ESC-derived iHPCs, which provides insights into B cell therapy.

## RESULTS

### Transplantation of iHPCs derived from *a Runx1-p2a-Hoxa9-t2a-Lhx2-*ESC line preferentially give rise to B lymphopoiesis in B-cell deficient mice

To induce B cell lymphopoiesis, we followed a two-step method of testing transcription factor combinations (Guo et al., 2020; Wang et al., 2020). An inducible expression cassette of *Runx1-p2a-Hoxa9-t2a-Lhx2* was introduced into the *Rosa26* locus of a GFP-transgenic mouse embryonic stem cell line (C57BL/6 background) to establish the *iR9X2*-ESC cell line (Figure S1A). Conditional expression of exogenous *Runx1, Hoxa9*, and *Lhx2* were confirmed in the presence of doxycycline (Figure S1B). Following the protocol of hematopoietic progenitor cells induction from ESC *in vitro* (Guo et al., 2020; Wang et al., 2020) (Figure 1A), BMP4 and VEGF were used to induce mesoderm differentiation and hemangioblast formation from embryoid body. AFT024-(mSCF/mIL3/mIL6/hFlt3L) cell line-cultured supernatants were used as conditioned medium (CM) for the *in vitro* induction of iHECs and subsequent iHPCs, as AFT024 CM is beneficial for the generation of iHPCs *in vitro* (Pereira et al., 2013). iHECs (CD31^+^CD41^+^CD45^−^c-Kit^+^CD201^+^) phenotypically resembling embryonic pre-HSCs (Zhou et al., 2016) were generated from *iR9X2*-ESC on day 6 to day 11 in the presence of doxycycline (Figure 1B). The iHECs co-cultured with OP9-DL1 feeder cells were further educated into Lin^-^c-Kit^+^Sca-1^+^ iHPCs on day11 to day21 in the presence of doxycycline (Figure 1C). To assess the engraftment potential of these iHPCs, we transplanted 5 million *iR9X2*-ESC-derived iHPCs (*iR9X2*-iHPCs) on day 21 into sublethally irradiated (5 Gy) B-cell-deficient μMT mice (*iR9X2*-μMT mice) with continuous doxycycline water feeding after transplantation. Four weeks after transplantation, we observed donor-derived GFP^+^CD45^+^CD19^+^ B cells, but no GFP^+^CD45^+^CD3/CD4/CD8^+^ T cells and no GFP^-^CD45^+^CD19^+^ B cells in the peripheral blood (PB) of *iR9X2*-μMT mice transplanted with the iHPCs (Figure 1D). We also observed donor-derived GFP^+^CD45^+^Mac1^+^ myeloid cells in the PB of recipients four weeks after transplantation (Figure 1D and Figure S1C). However, the donor-derived myeloid cells were transient and barely detectable in the PB of *iR9X2*-μMT mice at week 8 after transplantation (Figure S1C). Several independent experiments indicated that the engraftment rate of *iR9X2*-ESC derived iHPCs was 91.3 % (42/46 mice) and gave rise to a total average of 17.7% donor B cells in the PB of μMT recipients (n=46) at week 4 after transplantation (Figure 1E). Thus, inducible expression of *Runx1, Hoxa9*, and *Lhx2* leads ESC differentiation towards hematopoietic progenitors preferentially capable of B lymphopoiesis.

**Figure 1.**
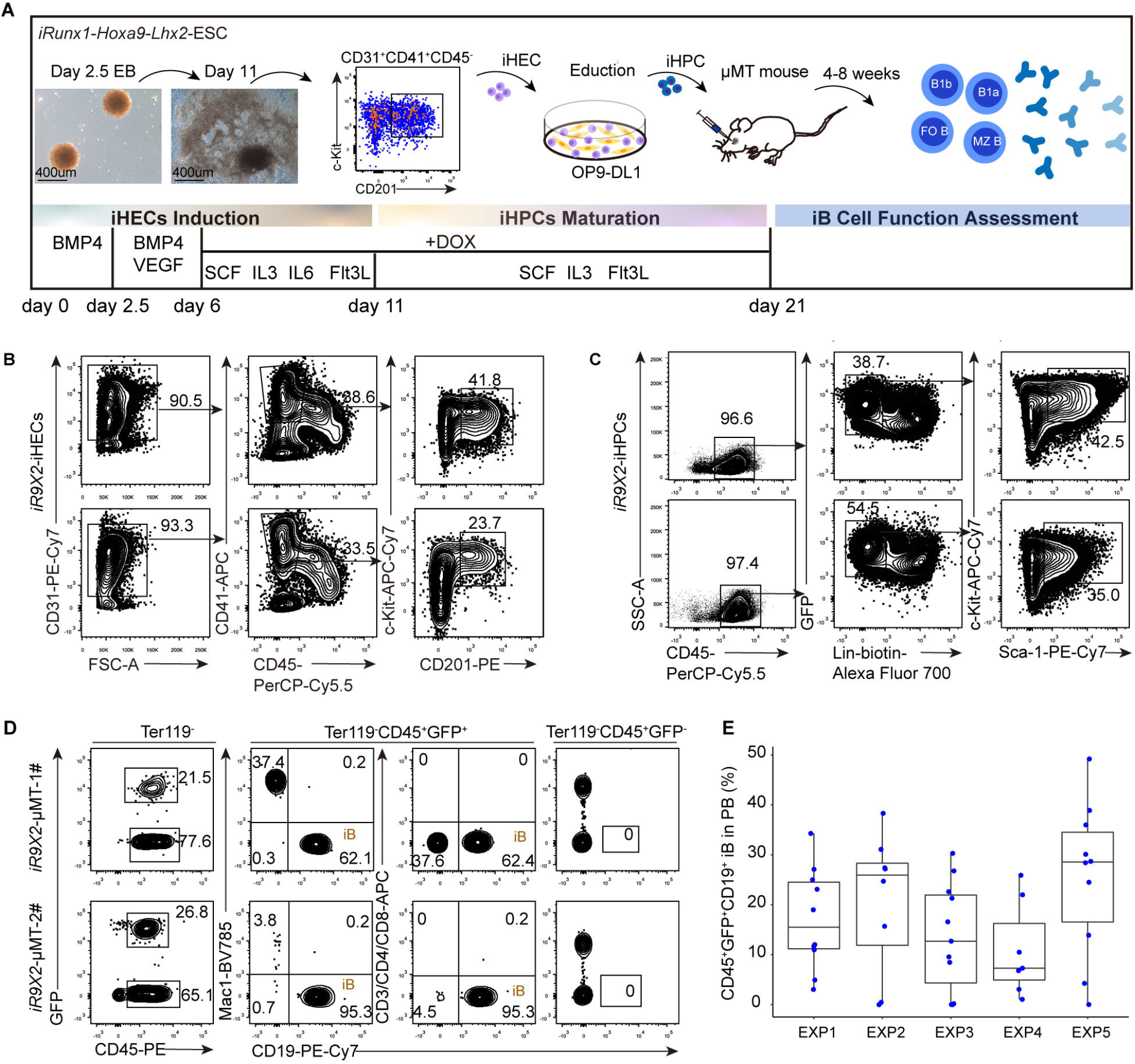
Reconstitution of B cells *in vivo* from *iRunx1-p2a-Hoxa9-t2a-Lhx2*-edited embryonic stem cells. (A) Schematic diagram of B cell regeneration from *iRunx1-p2a-Hoxa9-t2a-Lhx2* ESCs. ESC differentiation was initiated by embryoid body formation (EB). At day 2.5, EBs were replanted into gelatinized plates and differentiate to iHECs with cytokines. At day 11, iHECs were re-plated on OP9-DL1 stroma with cytokines for differentiation to hematopoietic progenitor cells. Inducible expression of *Runx1, Hoxa9*, and *Lhx2* were achieved by adding doxycycline (1 μg/mL, Sigma) during day 6 to day 21. At day 21, 5 million *iR9X2*-ESCs-derived iHPCs were injected into each sublethally irradiated (5 Gy) μMT mouse via retro-orbital veins. The mice were fed with water containing doxycycline (1 mg/mL) to induce the generation of B lymphocytes. After transplantation, B cell production was analyzed by flow cytometer and B cell function was evaluated. (B) Sorting strategies of iHECs population at day 11 induced from *iR9X2*-ESCs. The EB-derived iHECs (CD31^+^CD41^+^CD45^-^c-Kit^+^CD201^+^) were sorted by flow cytometry. Two representative plots from five independent experiments are shown. (C) Immuno-phenotypes of induced hematopoietic progenitor cells from iHECs after ten-day education. iHECs were co-cultured with OP9-DL1 for 10 days to generate iHPCs, and then fluorescence-activated cell sorting (FACS) analysis of iHPCs shows Lin^-^c-Kit^+^Sca-1^+^ phenotype. Two representative plots from five independent experiments are shown. Lin^−^ was defined as CD2^−^ CD3^−^ CD4^−^ CD8^−^ Mac1^−^ Gr1^−^ Ter119^−^ B220^−^ NK1.1^−^. (D) *iR9X2*-ESCs-derived B (iB) cells in the peripheral blood (PB) of μMT mice were analyzed by flow cytometry 4 weeks after transplantation. Five million iHECs-derived hematopoietic progenitors were transplanted into each sublethally irradiated μMT mouse (5 Gy). The mice were fed with water containing doxycycline (1 mg/mL) to induce the generation of B lymphocytes. iHPCs derived hematopoietic cells (GFP^+^CD45^+^Mac1^+^ myeloid cells, GFP^+^CD45^+^CD3/CD4/CD8^+^ T cells, and GFP^+^CD45^+^CD19^+^ B cells) were detected 4 weeks after transplantation. GFP^-^CD45^+^CD19^+^ gates of the same *iR9X2*-μMT recipients are shown as control. Two representative mice from five independent experiments are shown. (E) Summary of iB cells in the PB of individual μMT mice from five independent experiments. 46 μMT recipients were analyzed at week 4 after transplanting ESC-derived iHPCs. The box plot shows the percentage of the CD45^+^GFP^+^CD19^+^ iB cells in the PB. The percentage values were illustrated by ggplot2 (R package). One point represents one mouse. See also Figure S1.

To determine whether regenerative B (iB) cells possess antibody production ability, we quantified preimmune Ig isotype levels in the sera from *iR9X2*-μMT (iB-μMT) mice and μMT mice. We examined significant levels of serum IgM, IgG1, IgG2b, IgG2c, IgG3, and IgA in iB-μMT mice 4 to 6 weeks after iHPCs transplantation (Figure 2A), whereas basal levels of serum Ig isotypes in μMT mice could not be detected. 18 to 40 weeks after iHPCs transplantation, we could still detect significant preimmune Ig isotypes production levels in the sera from iB-μMT mice (Figure S2). The diversity of BCRs generated by the rearrangement of V(D)J gene segments in the B cell genomic locus (Tonegawa, 1983) is essential for humoral immune protection, as highly diverse antibody repertoires are capable of specifically binding a plethora of foreign antigens. To further assess the BCR repertoires of iB cells, we sorted naïve follicular B cells (FO B) (CD19^+^IgD^++^IgM^+^CD23^++^CD21^+^Lin^-^) for BCR deep sequencing from the spleen of one iB-μMT mouse at week 4 after transplantation and one C57BL/6 (B6) mouse (Figure 2B). The aliquots of 100,000 sorted naïve FO B were used as cell inputs for BCR sequencing at transcription level respectively. BCR clonotype profiling using MiXCR (Bolotin et al., 2015) captured abundant BCR sequences among the sorted naïve FO B cells isolated from the spleen of iB-μMT mouse that resembled their counterparts of B6 mouse (Figure 2C). Collectively, these data indicate that the humoral immune system is successfully reconstituted from *iR9X2*-ESC resource, with functional iB cells which express highly diverse BCR repertoires.

**Figure 2.**
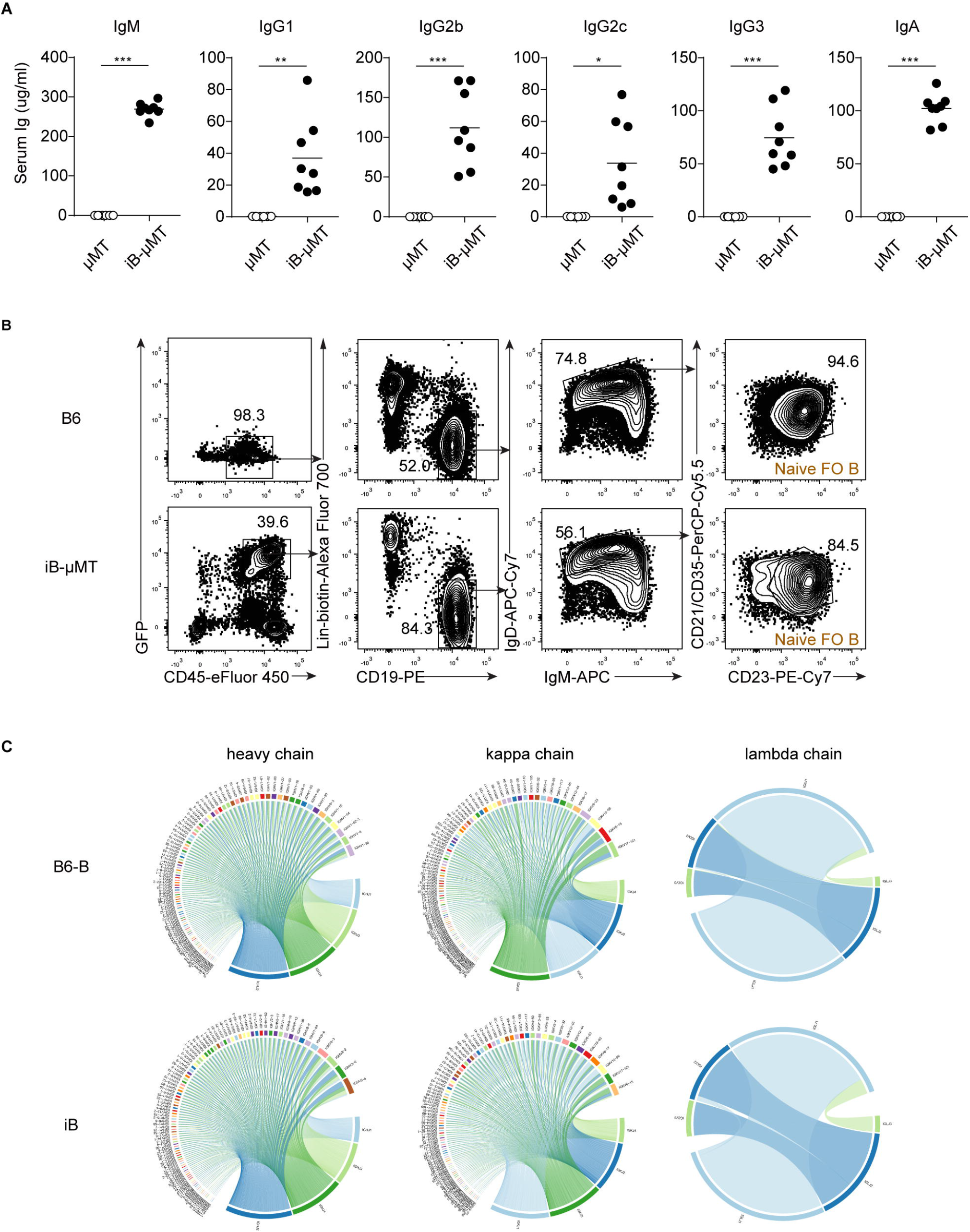
Basal levels of serum Ig in iB-μMT mice and the BCR repertoires of regenerative naïve FO B cells. (A) Serum Ig levels in iB-μMT mice and μMT mice (n=8 per group) were measured by ELISA. Sera were collected from iB-μMT mice (4 to 6 weeks after transplantation) and μMT mice. The different isotypes of antibodies (IgM /IgG1 /IgG2b /IgG2c /IgG3 /IgA) were measured by ELISA. Each symbol represents an individual mouse; small horizontal lines indicate the mean. *P < 0.05, **P < 0.01, and ***P < 0.001 (Independent-sample student t test). (B) Naïve follicular B (FO B) sorting strategy for BCR sequencing. One iB-μMT mouse was sacrificed 4 weeks after transplantation and one C57BL/6 (B6) mouse was sacrificed as control. From the spleen, naïve FO B cells were sorted based on CD19^+^ IgD^++^ IgM^+^ CD23^++^ CD21^+^ Lin^-^ (CD3^-^CD4^-^CD8^-^Ter119^-^Gr1^-^Mac1^-^NK1.1^-^CD138^-^) surface expression. (C) Chord diagram of IgH/IgK/IgL diversity in iB cells. Aliquots of sorted 100000 naïve FO B cells from the spleen of iB-μMT mouse or B6 mouse were used as cell inputs for BCR sequencing. The BCR IgH/IgL/IgK clonotypes were visualized in the form of chord diagram using VDJtools software(version 1.2.1). See also Figure S2.

### The regenerative B cell hierarchy shows a similar trajectory of natural B cell lymphopoiesis

We further analyzed the tissue distributions and immunophenotypes of the regenerative B lymphocytes in iB-μMT mice. We first detected induced pro-B cells and pre-B cells in the bone marrow where B lymphopoiesis starts. GFP^+^Lin^-^B220^+^CD43^+^ pro-B cells were detected in the bone marrow of iB-μMT mice at day 8 after *iR9X2*-iHPCs transplantation (Figure 3A). Induced pro-B cells could be separated into pre-pro-B (fraction A), early pro-B (fraction B), and late pro-B/early pre-B (fraction C/C′) cells according to Hardy’s criteria (Figure 3A) (Hardy et al., 1991). Then the induced pre-B cells which ceased CD43 expression and arose from pro-B cells appeared in the bone marrow of the same iB-μMT mice at day 14 after *iR9X2*-iHPCs transplantation (Figure 3A). Meanwhile, the majority of the induced pro-B cells were at early pro-B fraction at day 8 and further progressed into late pro-B/early pre-B fraction at day 14. Immature B cells which began to express complete surface IgM and mature B cells arose in the central bone marrow of iB-μMT mice at day 14 after *iR9X2*-iHPCs transplantation (Figure 3B). Then, GFP^+^CD45^+^CD93^+^B220^+^ transitional B cells which were early emigrant cells from the bone marrow were detected in the spleen of iB-μMT mice and could be divided into T1 population, T2 population, and T3 population according to the expression of surface IgM and CD23, and immature B cells further developed into GFP^+^CD45^+^CD93^-^B220^+^ mature B cells in the spleen of iB-μMT mice (Figure 3C). Interestingly, the majority of the iB cells in the spleen were transitional B cells at week 2 after transplantation, then further progressed into mature B cells at week 4 and week 8. Importantly, all mature B cell subsets of B-1a, B-1b, FO B, and MZ B existed in the spleen of iB-μMT mice at week 4 and week 8 after transplantation (Figure 3D). Induced B-1 and B-2 cells were also detected in the peritoneal cavity of iB-μMT mice at week 4 and week 8 after transplantation (Figure 3E). Furthermore, we could still observe all mature iB subsets of B-1a, B-1b, FO B, and MZ B in the spleen and peritoneum of iB-μMT mice at week 40 after transplantation (Figure S3A), despite that pro/pre-B and immature B cells were barely detected in the bone marrow of iB-μMT mice at week 6 after transplantation (Figure S3B), indicating a long life-span of mature iB cells. Taken together, these data indicate that the *iR9X2*-ESC-derived iHPCs reconstitute B lymphopoiesis *in vivo* in a spatiotemporal kinetic distribution pattern resembling natural B cell development.

**Figure 3.**
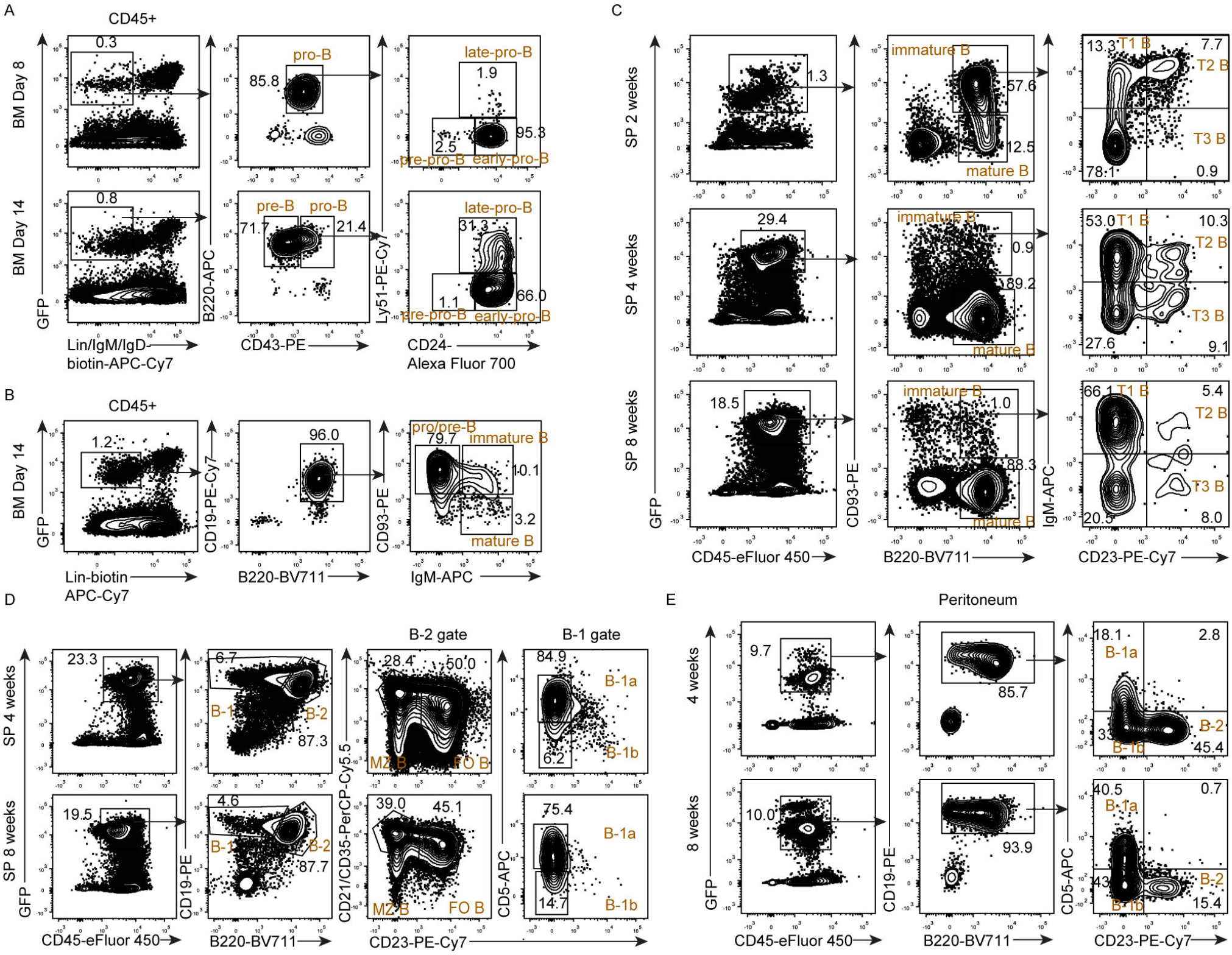
Cellular hierarchy of iB cell development in iB-μMT mice. (A) Flow cytometry analysis of pro-B cells and pre-B cells in the bone marrow of iB-μMT mice. Each μMT recipient was transplanted with five million *iR9X2*-iHPCs collected at day 21. The tibia of representative mouse was amputated and analyzed 8 days after transplantation; same mouse was sacrificed and analyzed 14 days after transplantation. Pro-B (GFP^+^Lin^-^IgM^-^IgD^-^B220^+^CD43^+^) and pre-B (GFP^+^Lin^-^IgM^-^IgD^-^B220^+^CD43^-^) from one representative mouse are shown. Pro-B cells are further divided into three subsets of pre-pro-B, early pro-B, and late pro-B on the basis of the expression of CD24 and Ly-51. Lin^-^ was defined as Ter119^-^Mac1^-^Gr1^-^NK1.1^-^CD3^-^CD4^-^CD8^-^. (B) Flow cytometry analysis of pro/pre-B cells, immature B cells, and mature B cells in the bone marrow of iB-μMT mice 14 days after transplantation. Each μMT recipient was transplanted with five million *iR9X2*-iHPCs collected at day 21. Data from one representative mouse are shown. Lin^-^ was defined as Ter119^-^ Mac1^-^Gr1^-^NK1.1^-^CD3^-^CD4^-^CD8^-^. (C) Transitional B cells were analyzed by flow cytometry from the spleen of iB-μMT mice. Each recipient was transplanted with five million *iR9X2*-iHPCs collected at day 21. iB-μMT mice were sacrificed and analyzed 2 weeks, 4 weeks, and 8 weeks after transplantation, respectively. Representative FACS plots from three iB-μMT mice are shown (D) Phenotypic analysis of B-1a, B-1b, follicular B (FO B), and marginal zone B (MZ B) cells in the spleen of iB-μMT mice 4 weeks and 8 weeks after transplantation. Each recipient was transplanted with five million *iR9X2*-iHPCs collected at day 21. Data from two representative mice are shown. (E) Phenotypic analysis of B-1a, B-1b, and B-2 cells in the peritoneal cavity of iB-μMT mice 4 weeks and 8 weeks after transplantation. Each recipient was transplanted with five million *iR9X2*-iHPCs collected at day 21. Data from two representative mice are shown. See also Figure S3.

### Transcriptome features of the regenerative pro-B and pre-B cells at single cell resolution

To characterize the transcriptome landscape of the early bone marrow regenerative progenitors in iB-μMT mice, we performed single-cell RNA-Seq using the sorted GFP^+^CD45^+^CD3^-^CD4^-^CD8^-^Ter119^-^Gr1^-^Mac1^-^NK1.1^-^ cells from the bone marrow of iB-μMT mice at day 7.5 after *iR9X2*-iHPCs transplantation (Figure S4A). Then the scRNA-seq datasets were processed and 4 clusters of a total of 7977 single cells including pro-B, large pre-B, megakaryocyte/erythrocyte progenitors (MEP), and granulocyte/macrophage progenitors (GMP) were identified and visualized using UMAP based on their unique gene expression signatures (Figure 4A). Two clusters were identified as B progenitors based on enrichment genes encoding proteins involved in B cell activation and B cell differentiation (Figure 4B and 4C), the surface marker-encoding gene *Cd19* (Figure 4D) which first takes place around the time of immunoglobulin gene rearrangement (Scheuermann and Racila, 1995), as well as the expression of *Cd93* (Figure 4D) which marks early B lineage cells (Li et al., 1996; McKearn et al., 1984). Pro-B cells (5754 single cells) were identified by surface marker-encoding genes including *Kit, Spn*, and *Cd24a*, while large pre-B cells (2013 single cells) were characterized by loss of *Kit* expression, occurrence of *Il2ra* expression and *Igkc* transcription, downregulated surface marker gene *Spn*, and upregulated surface marker gene *Cd24a* (Hardy et al., 1991; Osmond et al., 1998) (Figure 4D and 4E). During early B cell development, recombinase-activating genes (*Rag1/Rag2*) and DNA nucleotidylexotransferase (*Dntt*) were expressed (Figure S4B), which are essential for VDJ recombination (Bassing et al., 2002) at pro/pre-B cell stage. After immunoglobulin heavy chain rearrangement, the expression of *Dntt* was lost at large pre-B cells stage (Figure S4B) (Park and Osmond, 1987; Rolink et al., 1994), and the pre-BCR complex was formed by associating recombined heavy chain with VpreB and λ5 surrogate light chain (SLC) proteins (Figure 4E and S4B) and the Igα/CD79a and Igβ/CD79b signaling components (Figure 4E and S4B). The expression of pre-BCR at large pre-B cell stage is a critical early B cell development checkpoint for verifying functional production of μH. The signals from pre-BCR downregulated the expression of *Rag1/Rag2* (Figure S4B) (Grawunder et al., 1995), promoted the proliferation of large pre-B cells (Opstelten and Osmond, 1983; Osmond and Owen, 1984), and further silenced the expression of VpreB and λ5 (Figure 4E and S4B) (Burrows et al., 2002) for terminating large-pre B cells expansion and driving differentiating to small pre-B cell stage. Loss of any one component of pre-BCR complex will result in an arrest of B cell differentiation at the pro-B to pre-B cell stage (Gong and Nussenzweig, 1996; Kitamura et al., 1992; Kitamura et al., 1991; Martensson et al., 1999). Additionally, pro-B and large pre-B cells expressed the bruton’s tyrosine kinase (*Btk*) and B cell linker protein (*Blnk*) (Figure 4E), which are key cytoplasmic component genes of the pre-BCR signaling pathway (Kerner et al., 1995; Pappu et al., 1999).The transcriptional factor genes involved in regulatory network of early B cell development such as *Ikzf1, Spi1, Tcf3, Foxo1, Ebf1, Bcl11a*, and *Pax5*, were widely expressed among pro-B and large pre-B population (Figure 4F and S4B), and loss of any one will result in an arrest of B cell differentiation (Reynaud et al., 2008; Scott et al., 1994; Bain et al., 1994; Dengler et al., 2008; Lin and Grosschedl, 1995; Yu et al., 2012b; Urbanek et al., 1994). Also, pro-B cells showed abundant expression of transcription factor *Erg* which was reduced at large pre-B stage (Figure 4F), suggesting an exquisitely stage-specific regulator of early B-cell development (Ng et al., 2020). While large pre-B cells widely expressed transcription factor *Bach2* (Figure 4F), which is required for mediating negative selection at the pre-BCR checkpoint (Swaminathan et al., 2013). And the expression of transcription factors *Irf4* and *Ikzf3* were upregulated by pre-BCR signaling (Muljo and Schlissel, 2003; Thompson et al., 2007) at large pre-B cell stage (Figure 4F), which will play a critical role in further downregulating pre-BCR and suppressing large pre-B cells expansion for transition from large pre-B to small pre-B cells (Ma et al., 2010; Ma et al., 2008). In addition, two small clusters of MEP cells and GMP cells (141 single cells and 69 single cells respectively) were marked by high expression of carboxylate reductases (*Car1* and *Car2*) (Villeval et al., 1985) and a number of granule genes including myeloperoxidase (*Mpo*), neutrophil elastase (*Elane*), proteinase 3 (*Prtn3*), and cathepsin G (*Ctsg*) respectively (Figure 4B), which addresses the transient wave of myeloid lineage cells in the iB-μMT mice. Together, large-scale single cell transcriptome features demonstrate that *iR9X2*-iHPCs robustly differentiate into early B cell progenitors as early as day 7.5 after transplantation.

**Figure 4.**
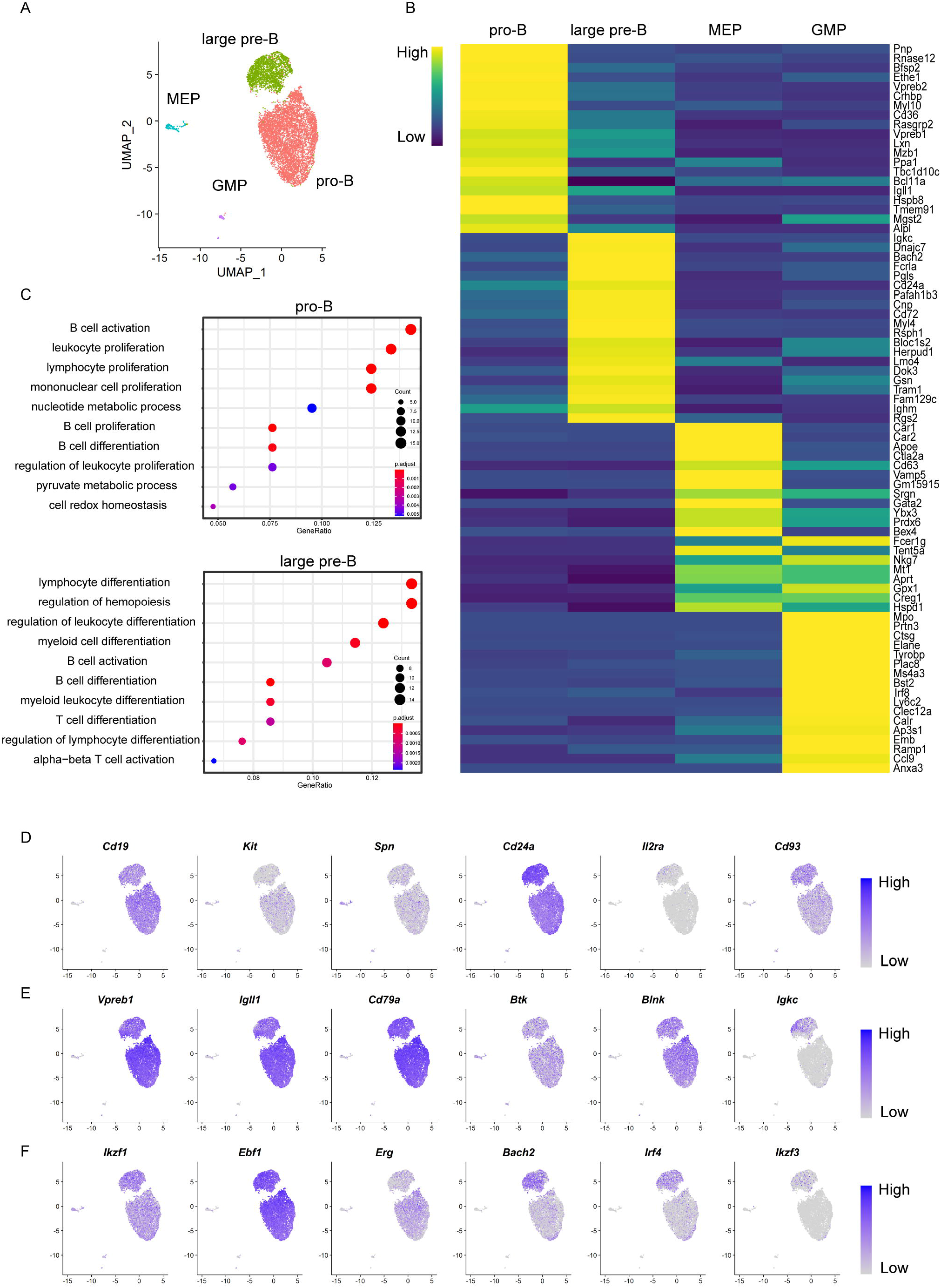
Single-cell transcriptomic characterization of regenerative pro-B and pre-B cells. (A) UMAP visualization of early bone marrow regenerative progenitor cells in iB-μMT mice. 50000 bone marrow cells were sorted based on GFP^+^CD45^+^CD3^-^CD4^-^CD8^-^Ter119^-^Gr1^-^Mac1^-^NK1.1^-^ surface expression for sequencing at day7.5 after transplantation. 7977 cells were retained for UMAP analysis. To rule out the effects of cell cycle variances, we performed a simple linear regression against the cell cycle score calculated by CellCycleScoring. Then, PCA was performed by RunPCA using 2000 highly variable genes and the top 20 PCs were used for UMAP analysis. Clusters were detected using FindClusters with setting parameters dims = 1:20, resolution = 0.08. One point represent one cell. (B) Heatmap showing the average expression of top 20 differential expressed genes in each clusters of early bone marrow regenerative progenitor cells from iB-μMT mice. Up-regulation genes were identified for each cluster using Wilcoxon rank sum test with parameter ‘min.pct = 0.5, logfc.threshold = 0.25’ implemented in Seurat. One cluster per column. The expression value were z-score transformed by Seurat3 packages. Heatmap were plotted by pheatmap (version 1.0.12). (C) Gene ontology (GO)–enrichment analysis of the differentially expressed genes in pro-B and large pre-B clusters. Gene ontology-enrichment analysis (biological processes) were performed for the up-regulation genes of each cluster by clusterProfiler with BH adjusted p-value cutoff =0.05 (version 3.14.3) (Yu et al., 2012a). Each symbol represents a GO term (noted in plot); color indicates adjusted P value (Padj (significance of the GO term); bottom key), and symbol size is proportional to the number of genes (top key). (D) UMAP analysis of the expression patterns of pro-B cells and large pre-B cells related surface marker genes (*CD19, Kit, Spn, Cd24a, Il2ra, Cd93*). The expression value of each gene was converted by log2 and illustrated by FeaturePlots of Seurat3. (E) UMAP analysis of the expression patterns of selected pre-BCR and BCR formation related marker genes (*Vpreb1, Igll1, CD79a, Btk, Blnk, Igkc*). The expression value of each gene was converted by log2 and illustrated by FeaturePlots of Seurat3. (F) UMAP analysis of the expression patterns of selected early B cell development related transcription factors (*Ikzf1, Ebf1, Erg, Bach2, Irf4, Ikzf3*). The expression value of each gene was converted by log2 and illustrated by FeaturePlots of Seurat3 See also Figure S4.

### The regenerative B cells produce adaptive immune response and form long-term immune memory

To investigate the immune function of regenerative B cells, we performed inoculation of T cell dependent antigen (TD Ag) to test the humoral immune response in iB-μMT mice. We immunized iB-μMT mice with 4-hydroxy-3-nitrophenylacetyl-chicken-gammma-globulin conjugates (NP-CGG) and detected the levels of NP-specific IgM and IgG1 antibodies in the sera from immunized mice (Figure 5A). The iB-μMT mice showed elevated NP-specific IgM, total NP-specific IgG1, and high affinity NP-specific IgG1 levels compared with μMT mice after primary immune response (Figure 5B). After boosting with NP-CGG, increased amounts of total and high-affinity NP-specific IgG1 antibodies were produced quickly from iB-μMT mice while antibodies were not detected in μMT mice (Figure 5C). We next assessed the normal formation of germinal centers (GC) B cells and the generation of memory B cells and plasma cells in iB-μMT mice which adaptive humoral immune protection relies heavily on. Two weeks after NP-CGG immunization, there were robust emergence of plasma cells (B220^low/-^CD138^+^) and NP-specific GC B cells (NP^+^GL7^+^CD38^-^) in the spleen of iB-μMT mice which were comparable to the B6 mice counterparts (Figure 6A). In addition, we could detect antigen specific class-switched IgG1^+^ memory B cells in the spleen of iB-μMT mice at day 21 after NP-CGG immunization (Figure 6B), suggesting a successful process of immunoglobulin class switching. And abundant long lived plasma cells (Lin^-^IgM^-^CD22^-^CD19^-^MHCII^-^CD138^+^) in the bone marrow of iB-μMT mice 3 weeks after NP-CGG immunization were detected, which was comparable to the B6 mice counterpart (Figure 6C). Importantly, long lived plasma cells could still be detected in the bone marrow of iB-μMT mice at day 17 after the boost (Figure 6C). Thus, these results indicate that the regenerative iB cells in iB-μMT mice could produce primary response and memory response and sustain a long-term humoral immunity protection, suggestive of a typical adaptive immune response.

**Figure 5.**
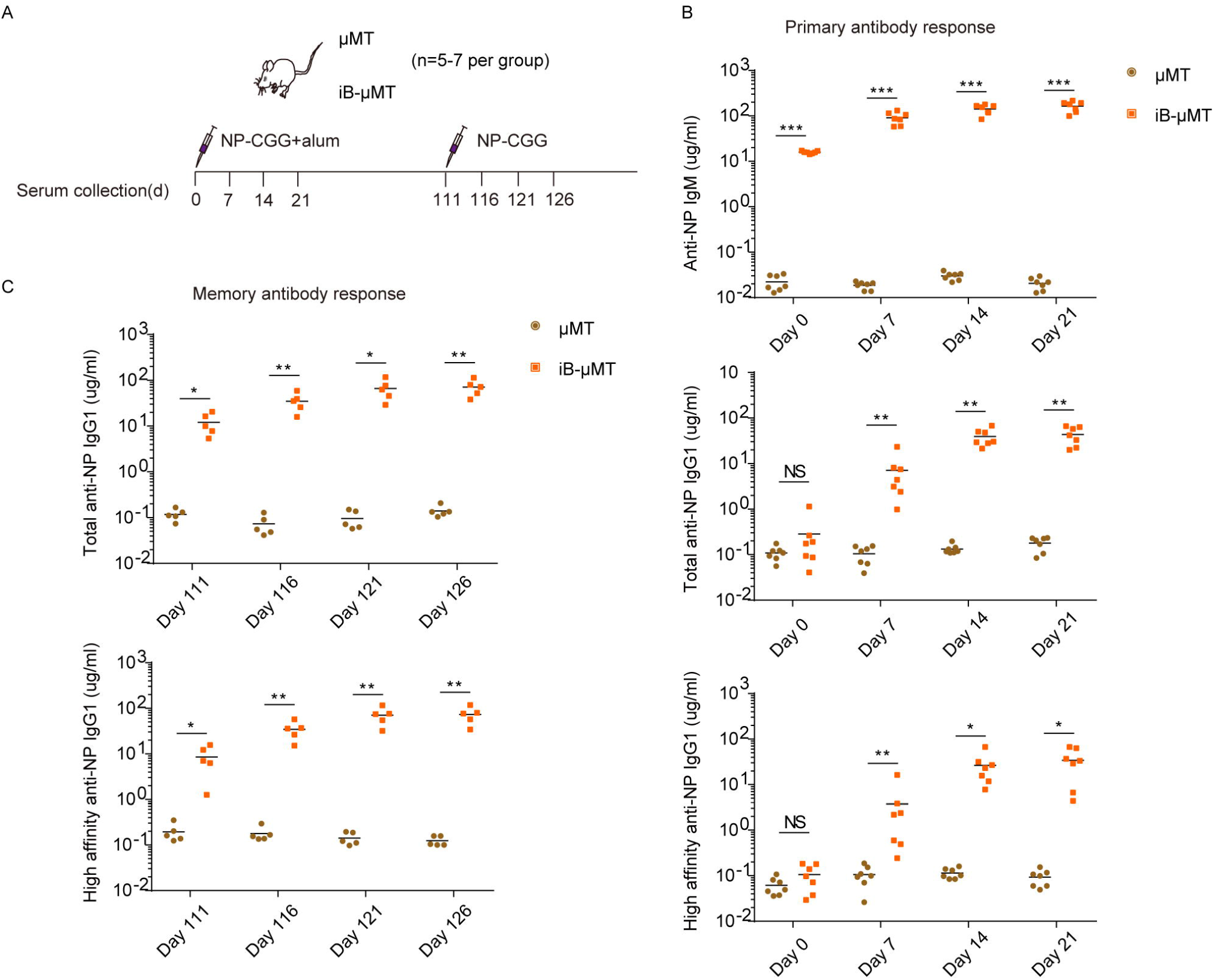
Antibody response to T-cell-dependent antigen (NP-CGG) in iB-μMT mice. (A) Schematic diagram of iB-μMT mice and μMT mice immunization with NP-CGG and sera collection at indicated time points. iB-μMT mice (n=7) and μMT mice (n=7) were given primary intraperitoneal immunization with 100μg NP-CGG in alum at day 0. Then iB-μMT mice (n=5) and μMT mice (n=5) were given secondary challenge with 50μg NP-CGG at day 111 after primary immunization. Sera were collected at day 0, day 7, day 14, day 21, day 111, day 116, day 121, and day 126 after primary immunization. (B) T-cell-dependent primary antibody response in iB-μMT mice. iB-μMT mice (n=7) and μMT mice (n=7) were immunized with 100μg NP-CGG in alum at day 0. Anti-NP IgM (top), total anti-NP IgG1 (middle), and high-affinity anti-NP IgG1 (bottom) antibodies in the sera collected at indicated time points were measured by ELISA. Each symbol represents an individual mouse, the horizontal lines indicate the mean values. NS, not significant, *P < 0.05, **P < 0.01, and ***P < 0.001 (Independent-sample student t test and Mann-Whitney U test). (C) Production of memory antibodies in iB-μMT mice. Sera were collected from iB-μMT mice (n=5) and μMT mice (n=5) at indicated time points after boosting with 50μg NP-CGG. Levels of total anti-NP IgG1 and high-affinity anti-NP IgG1 were measured by ELISA. Each symbol represents a mouse, the horizontal lines indicate the mean values. *P < 0.05, **P < 0.01, and ***P < 0.001 (Independent-sample student t test and Mann-Whitney U test).

**Figure 6.**
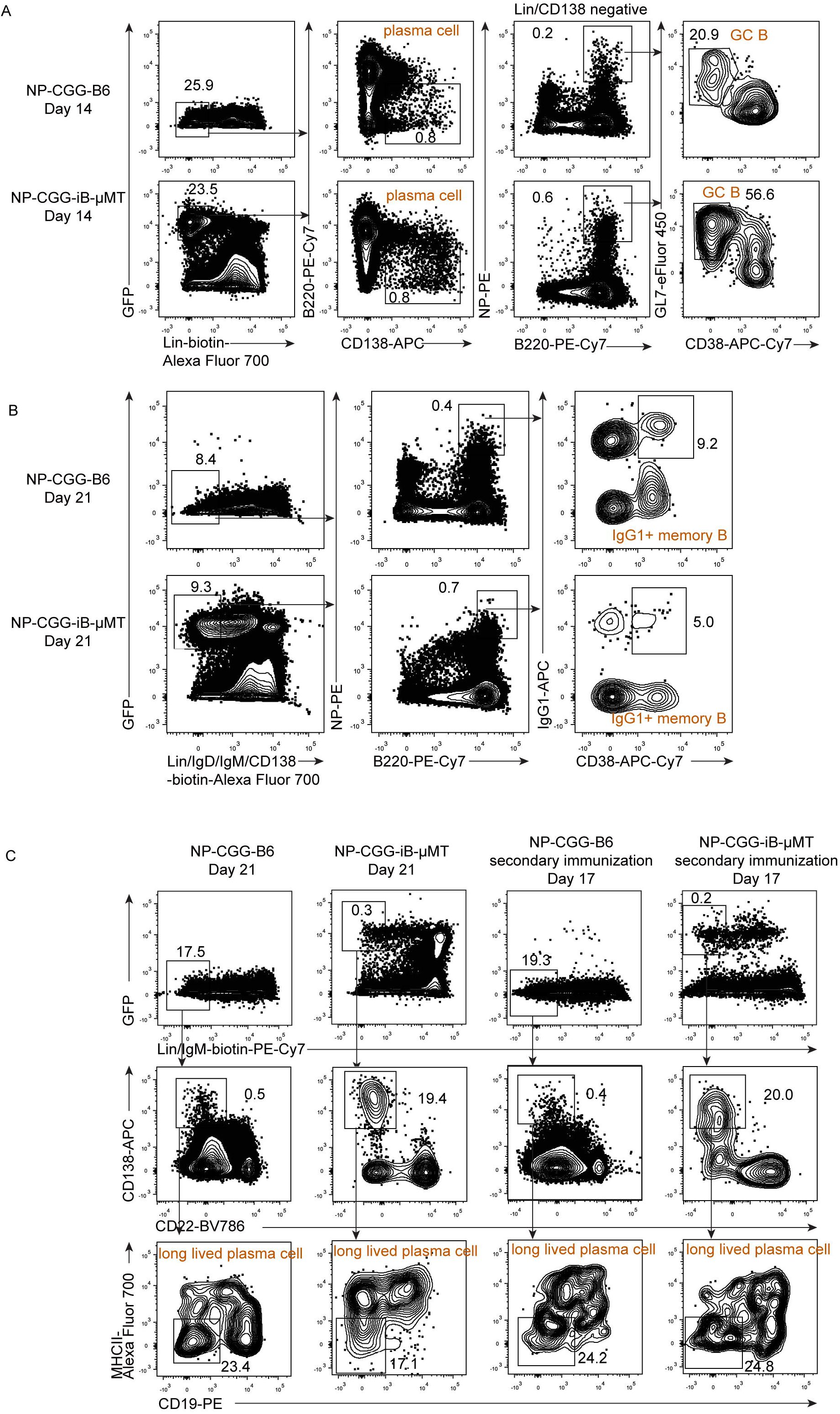
Normal germinal center (GC) B, memory B, and plasma cell formation in iB*-*μMT mice. (A) Plasma cells and antigen-specific GC B cells in the spleen of iB-μMT mice. Spleen cells were isolated from B6 and iB-μMT mice 14 days after primary intraperitoneal immunization with 100μg NP-CGG in alum. Plasma cells (Lin^-^B220^low/-^CD138^+^) and antigen-specific GC B cells (Lin^-^B220^+^CD138^-^NP^+^GL7^+^CD38^-^) were analyzed by flow cytometry. Data from one representative B6 mouse and one representative iB-μMT mouse are shown. Lin^-^ was defined as Ter119^−^Mac1^−^Gr1^−^NK1.1^−^CD3^−^CD4^-^CD8^-^. (B) Antigen-specific class-switched IgG1^+^ memory B cells in the spleen of iB-μMT mice. Spleen cells were isolated from B6 and iB-μMT mice 21 days after primary intraperitoneal immunization with 100μg NP-CGG in alum. Representative flow cytometry plots of antigen-specific class switched IgG1^+^ memory B cells (Lin^-^IgD^-^IgM^-^CD138^-^B220^+^NP^+^CD38^+^IgG1^+^) from one B6 mouse and one iB-μMT mouse are shown. Lin^-^ was defined as Ter119^−^ Mac1^−^Gr1^−^NK1.1^−^CD3^−^CD4^-^CD8^-^. (C) Long lived plasma cells in the bone marrow of iB-μMT mice. Long lived plasma cells (Lin^-^IgM^-^CD138^+^CD22^-^CD19^-^MHCII^-^) in the bone marrow of B6 and iB-μMT mice were analyzed by flow cytometry. Representative plots from B6 and iB-μMT mice 21 days after priming with NP-CGG in alum and 17 days after NP-CGG boosting are shown. Lin^-^ was defined as Ter119^−^Mac1^−^Gr1^−^NK1.1^−^CD3^−^CD4^-^CD8^-^.

## DISCUSSION

In this study, we demonstrated that enforced expression of three transcription factor *Runx1, Hoxa9*, and *Lhx2*, can guide an abundant B lineage fate commitment and *in vivo* B lymphopoiesis in B-cell deficient animals. Of note, the abundancy of *in vivo* lymphopoiesis includes pro/pre-B progenitors, immature B cells, and all subsets of mature B-1a, B-1b, FO B, and MZ B cells. This HSC-independent approach regenerates a complete humoral system which rescues B cell-related immune responses in the inherited B-cell deficient animals.

We still detected the presence of induced mature B-1 and B-2 cells, and serum antibodies in iB-μMT mice 40 weeks after transplantation. Transplantation of mouse ESC differentiated B220^+^CD93^+^ pro/pre-B cells reconstituted B-1 and B-2 cells in recipients, however the B cell regeneration was transient and the serum contained only low to undetectable IgM antibodies, which were barley detected 6-8 weeks after transplantation (Potocnik et al., 1997). We noticed that in our system the iHPCs in the bone marrow microenvironment differentiated into pro/pre-B cells and further matured into B cells *in vivo* might avoid the partial failure of microenvironment-modeling of B cell development *in vitro* (Potocnik et al., 1997). Constitutive expression of *Lhx2* in natural hematopoietic progenitor/stem cells *in vivo* developed a myeloproliferative disorder and caused acute leukemia (Richter et al., 2003), which implies that the iHPCs generated in our study are different from these cells as our iHPCs have never caused myeloid proliferation. Certain pre-B tumor cell lines expressed *Lhx2* (Xu et al., 1993) but in our case we have never observed B cell tumors, indicating that synergistic expression of *Lhx2* with *Runx1* and *Hoxa9* starting from an early stage of hematopoietic development prior to definitive HSC occurrence leads to no obvious tumorigenic effect. In addition, single-cell RNA-seq showed typical expression patterns of surface markers, transcription factors and essential regulators, and pre-BCR complex in induced pro-B and large pre-B that suggests a normal early B lymphopoiesis in iB-μMT mice. Of note, *Ikzf1, Spi1, Tcf3*, and *Pax5* are normally expressed in the induced B cell progenitors, as reduction or loss of these master factors are associated with B cell leukemia (Hu et al., 2016; Okuno and Yuki, 2012; Aspland et al., 2001; Liu et al., 2014).

Synergistic expression of *Runx1* and *Hoxa9* during mESC differentiation resulted in a iHPC population that preferentially contributes to T lymphopoiesis *in vivo* (Guo et al., 2020). In this study, using the same induction protocol, coordination of *Runx1, Hoxa9*, and *Lhx2* promotes B lymphopoiesis instead of T lymphopoiesis *in vivo*. It has been reported that T cell development was blocked by the expression of *Lhx2 in vivo* in HSPC (Kitajima et al., 2015). Surprisingly, synergistic expression of *Runx1* and *Lhx2* transcription factors preferentially determined T cell lineage fate using the same induction system (Figure S5), despite the low efficiency compared with the effect of *Runx1* and *Hoxa9*. Thus our data demonstrate that the synergistic effects of *Runx1, Hoxa9*, and *Lhx2* transcription factors are more complicated than a simple addition-subtraction methodology. It is worth of further investigation to comprehensively understand the epigenetic landscape architected by *Runx1, Hoxa9*, and *Lhx2* during hematopoietic fate commitment and subsequent B lymphopoiesis.

In conclusion, this study establishes a novel approach of reconstituting complete B lymphopoiesis *in vivo* based on a two-step approach of in vitro HPC commitment from PSC and *in vivo* lymphopoiesis. The regenerative B cells possess abundant BCR repertoires capable of recognizing numerous different antigens and can restore adaptive humoral immune response and form immune memory in B cell-deficient mice. The robust induction system of B cell regeneration provides a new tool for the basic study of B cell development and B cell disease modeling. Given the unlimited cell resource and gene-editing advantages of PSCs, our study provides an insight into therapeutic applications of regenerative B cells for individuals suffering from inherited B cell defects.

### Limitations of the study

We have identified that the synergistic expression of *Runx1, Hoxa9*, and *Lhx2* directs B lymphopoiesis from PSC, while determining the underlying mechanisms of these transcription factors during B cell lineage fate commitment will be a critical goal. And, human B cell regeneration using human PSCs source by synergistic expression of these or other transcription factors is the key objective of future study, which will provide insights into B-cell immunotherapy and vaccine study.

## ACKNOWLEDGMENTS

This work was supported by the National Key R&D Program of China (2019YFA0110203, 2020YFA0112404), the Strategic Priority Research Program of the Chinese Academy of Sciences (XDA16010601), the Key R&D Program of Guangdong Province (2020B1111470001), the National Natural Science Foundation of China (81925002), the Key Research & Development Program of Guangzhou Regenerative Medicine and Health Guangdong Laboratory (2018GZR110104006), the Frontier Science Research Program of the CAS (QYZDB-SSW-SMC057), and the Science and Technology Planning Project of Guangdong Province, China (2020B1212060052).

## AUTHOR CONTRIBUTIONS

Conceptualization, J.W., Q.Z., and B.W.; Methodology, validation, and analysis, Q.Z. and B.W.; BCR sequencing and scRNA-seq data analysis, Q.W. and Y.L.; Investigation, Q.Z., B.W., F.H., C.X., H.P., Y.W., X.L., L.L., J.X., and Y.Z.; Resources, Q.Z., B.W., X.L., L.L., Y.G., and J.W.; Writing – original draft, J.W. and Q.Z.; Writing – review & editing and visualization, J.W., Q.Z., B.W., Q.W., J.D., and M.Z.; Funding acquisition, J.W. and J.D.; Supervision, J.W.

## STAR METHODS

## RESOURCE AVAILABILITY

### Lead contact

Further information and requests for resources and reagents should be directed to and will be fulfilled by the lead contact, Jinyong Wang.

### Materials availability

This study did not generate new unique reagents.

### Data and code availability

The BCR sequencing and scRNA-seq datasets generated in this study are available on GEO: GSE180318 and GSE180319.

## EXPERIMENTAL MODEL AND SUBJECT DETAILS

### Animal used in this study

μMT (B6.129S2-Ighmtm1Cgn/J, CD45.2^+^) mice were purchased from The Jackson Laboratory. C57BL/6 (CD45.2^+^) mice were purchased from Bejing Vital River Laboratory Animal Technology. *Rag1*^−/−^mice (C57BL/6 background) were a gift from Dr. Zhihua Liu from the Institute of Biophysics (CAS, China). Mice were housed in the SPF-grade animal facility of the Guangzhou Institutes of Biomedicine and Health, Chinese Academy of Sciences (GIBH, CAS, China). All animal experiments were approved by the Institutional Animal Care and Use Committee of Guangzhou Institutes of Biomedicine and Health (IACUC-GIBH).

## METHOD DETAILS

### Cell culture

Mouse embryonic fibroblasts (MEFs) were derived from 13.5 d.p.c C57BL/6 mouse embryos. MEFs were maintained in DMEM/High glucose (Hyclone), 10% FBS (Natocor) supplemented with 1% nonessential amino acids (NEAA, Gibco). C57BL/6 mouse embryonic stem cells (Biocytogen) were maintained on feeder layers in ES medium containing DMEM/high glucose, 15% FBS (Gibco), 1% NEAA (Gibco), 1% GlutaMAX (Gibco), 1% Sodium Pyruvate (Gibco), 0.1 mM β-mercaptoethanol (Sigma), 1 μM PD0325901 (Selleck), 3 μM CHIR-99021 (Selleck), and 1000 U/mL LIF (Peprotech). The OP9-DL1 cells (GFP^+^) were maintained in α-MEM (Gibco) supplemented with 20% FBS (Ausbian). The AFT024 cell lines (ATCC) were maintained in DMEM/ high glucose, 10% FBS (Natocor) supplemented with 0.1 mM β-mercaptoethanol and 1% Sodium Pyruvate.

### Hematopoietic differentiation

ESCs were trypsinized by 0.05% Trypsin-EDTA (Gibco) and resuspended in the basic differentiation medium (BDM: IMDM, 15% FBS (Gibco), 200 μg/mL iron-saturated transferrin (Sigma), 0.1 mM β-mercaptoethanol (Sigma), 1% GlutaMAX, and 50 μg/mL ascorbic acid (Sigma)). To remove the feeder layers, the PSCs were plated into the 0.1% gelatin-coated (Merck Millipore) well, and the floating cells were collected after 30 min. For embryoid body (EB) generation, the PSCs were resuspended at 100,000 cells/mL in the BDM supplemented with 5 ng/mL BMP4 (Peprotech) and plated at 20 uL/drop for inverted culture in 15 cm dishes. At day 2.5, EBs were replanted into gelatinized plates in BDM supplemented with 5 ng/mL BMP4 and 5 ng/mL VEGF (Novoprotein). At day 6, the medium was changed to BDM supplemented with 2% conditioned medium derived from the supernatants of AFT024-mIL3, AFT024-mIL6, AFT024-hFlt3L, and AFT024-mSCF cell culture. Doxycycline (1 μg/mL, Sigma) was added at day 6. The medium was replaced every other day. The plates were seeded with OP9-DL1 cells (20,000 cells/well, 12-well plate) 12 h prior to the hematopoietic maturation step in EM medium (α-MEM, 15% FBS (Hyclone), 200 μg/mL iron-saturated transferrin, 0.1 mM β-mercaptoethanol, 1% GlutaMAX, 50 μg/mL ascorbic acid, 2% conditioned medium derived from supernatants of AFT024-mIL3, AFT024-hFlt3L, and AFT024-mSCF cell culture and 1 μg/mL doxycycline). Then, 1000–3000 sorted iHECs were seeded into each well for hematopoietic maturation. The EM medium was half replaced every two days.

### Transplantation of iHPCs

Eight-ten-week-old μMT mice and *Rag1*^−/−^mice were sublethally irradiated (5 Gy and 3.5 Gy respectively) by an X-ray irradiator (RS2000, Rad Source Inc.). 5 million *iR9X2*-ESC-derived iHPCs were injected into each irradiated μMT mouse via retro-orbital veins. And 3 million *iRunx1-Lhx2*-ESC-derived iHPCs were injected into each irradiated *Rag1*^*-/-*^ mouse via retro-orbital veins. The mice were fed with water containing doxycycline (1 mg/mL) to induce the generation of B lymphocytes.

### Gene editing

To generate GFP positive ESCs (GFP-ESC), the *CAG Pr-GFP-PGK Pr-PuroR* cassette was inserted into the *Hipp11* locus of mouse ESC (C57BL/6 background, CD45.2 strain). The positive clones (GFP-ESC) were selected by puromycine (1 μg/ mL, Thermo Fischer Scientific), and the expression of GFP was confirmed by flow cytometry. To generate *iRunx1-Hoxa9-Lhx2* (*iR9X2*) ESCs, a *CAG Pr-rtTA-3 × Stop-TRE-Runx1-p2a-Hoxa9-t2a-Lhx2-pA-PGK Pr-HygroR* cassette was inserted into the *Rosa26* locus of GFP-ESC. The positive clones (*iR9X2*-ESC) selected by hygromycin B (150 μg/ mL, Invivogen) were further cultured in ES medium supplemented with doxycycline (1 μg/mL, Sigma), and the induced expression of *Runx1, Hoxa9*, and *Lhx2* were confirmed by qPCR. And *a CAG Pr-rtTA-3 × Stop-TRE-Runx1-p2a-Lhx2-PGK Pr-HygroR* cassette was inserted into the *Rosa26* locus of GFP-ESC for generating *iRunx1-Lhx2* ESCs, the positive clones (*iRunx1-Lhx2*-ESC) selected by hygromycin B (150 μg/ mL, Invivogen) were further cultured in ES medium supplemented with doxycycline (1 μg/mL, Sigma), and the induced expression of *Runx1* and *Lhx2* were confirmed by qPCR.

### Flow cytometry and cell sorting

Single-cell suspensions were prepared in phosphate-buffered saline (PBS) supplemented with 2% fetal bovine serum (FBS) and filtered by 70 μm filter. Single cells were blocked by Fc (CD16/32) (Biolegend) antibody, and then stained with related antibodies. The following antibodies were used: c-Kit (2B8, eBioscience), CD31 (390, eBioscience), CD41 (eBioMWReg30, eBioscience), CD45 (30-F11, eBioscience), CD201 (eBio1560, eBioscience), CD2 (RM2-5, eBioscience), CD3 (145-2C11, eBioscience), CD4 (GK1.5, eBioscience), CD8a (536.7, eBioscience), B220 (RA3-6B2, eBioscience), B220 (RA3-6B2, BioLegend), Mac1 (M1/70, eBioscience), Mac1 (M1/70, BioLegend), NK1.1 (PK136, BioLegend), NK1.1 (PK136, eBioscience), Ter119 (TER-119, eBioscience), Gr1 (RB6-8C5, eBioscience), IgM (II/41, eBioscience), IgD (11-26c.2a, BioLegend), Sca-1 (D7, eBioscience), CD19(eBio1D3, eBioscience), CD23(B3B4, BioLegend), CD21/35(7G6,BD Biosciences), CD43(eBioR2/60, eBioscience), CD24(M1/69, BioLegend), Ly-51(6C3, BioLegend), CD93(AA4.1, BioLegend), CD5(53-7.3, BioLegend), CD138(281-2, BioLegend), CD38(90, BioLegend), GL7 (GL-7, eBioscience), IgG1(RMG1-1, BioLegend), CD22(Cy34.1, BD Biosciences), MHC II (M5/114.15.2, BioLegend), Streptavidin Alexa Fluor® 700 (Invitrogen), Streptavidin PE-Cy7 (BioLegend), NP-PE(Biosearch Technologies). The cells were resuspended in the DAPI solution (Sigma), or PI solution (BioLegend) and were analyzed with Fortessa cytometer (BD Biosciences). The cells were sorted using Arial III cytometer (BD Biosciences). The flow cytometry data were analyzed with FlowJo.

### Immunization and serum collection

T-cell–dependent antigen immunization was performed as described previously (Dai et al., 2007; Zheng et al., 2013). Briefly, iB-μMT mice 4 weeks after transplantation and μMT mice were immunized i.p. with 100 μg 4-hydroxy-3-nitrophenyl acetyl (NP)-CGG (Biosearch Technologies) in alum (Thermo Fisher Scientific), at volume ratio of 1:1 (200 μl/mouse). For recall responses, mice were challenged with 50 μg NP-CGG at week 16 after primary immunization (100 μl/mouse). Sera were collected from each group at day 0, day7, day 14, day 21, day 111, day 116, day 121, and day126 after primary immunization. Antigen-specific antibodies were measured by ELISA.

### ELISA

For basal serum Abs (IgM /IgG1 /IgG2b /IgG2c /IgG3 /IgA) measurement, microtiter plates were coated with Goat Anti-Mouse Ig (5μg/ml, Southern Biotech) overnight at 4 °C. For NP-specific Abs measurement, NP(27)-BSA (Biosearch Technology) or NP(9)-BSA (for high affinity) (Biosearch Technology) was used as the capture antigen. Then nonspecific binding was blocked with 0.5% BSA in PBS for 2 hours at 37 °C. Diluted serum samples were incubated in plates for 1 hours at 37 °C. Plates were incubated for 1 h with Goat Anti-Mouse IgA-HRP, Goat Anti-Mouse IgM-HRP, Goat Anti-Mouse IgG1-HRP, Goat Anti-Mouse IgG2b-HRP, Goat Anti-Mouse IgG2c-HRP, Goat Anti-Mouse IgG3-HRP (all from Southern Biotech), and then for 15-30min with 100ul/well TMB (BioLegend) substrate solution, followed by 50ul 2N H2SO4 to stop reaction. Absorbance values were read at 450nm using a microplate reader (Cytation5, BioTek).

### BCR sequencing

For BCR sequencing, 100000 naïve FO B were sorted from the spleen of one iB-μMT mouse 4 weeks after transplantation and one C57BL/6 (B6) mouse respectively. The sorted naïve FO B cells were gated on CD45^+^CD19^+^IgD^++^IgM^+^CD23^++^CD21^+^CD3^-^CD4^-^CD8^-^Ter119^-^Gr1^-^Mac1^-^NK1.1^-^CD138^-^. Total RNA was extracted from the naïve FO B cells using Trizol (MRC). 5’ RACE was performed with SMARTer RACE cDNA Amplification Kit (Clontech). IgG /IgK/IgL NGS libraries were made by using NEBNext Ultra DNA Library Prep Kit for Illumina (NEB). Libraries were sequenced on the Illumina Miseq 2 × 300 platform. The raw data (fastq files) were generated using illumina bcl2fastq software and were uploaded to Gene Expression Omnibus public database. The B cell receptor repertoires were aligned and assembled using software MiXCR (version 3.0.13). The BCR IgH/IgL/IgK clonotypes were exported respectively by parameter ‘--chains’ in exportClones command of MiXCR (Bolotin et al., 2015). The exported clonotypes were visualized in the form of chord diagram using VDJtools software (version 1.2.1) (Shugay et al., 2015).

### scRNA-seq and data analysis

50000 sorted early bone marrow regenerative progenitors (GFP^+^CD45^+^CD3^-^CD4^-^CD8^-^Ter119^-^Gr1^-^Mac1^-^NK1.1^-^) from iB-μMT mice (n=4) at day 7.5 after transplantation were used for scRNA-seq. Droplet-based scRNA-seq datasets were produced using a Chromium system (10x Genomics, PN120263) following manufacture’s instruction. Droplet-based scRNA-seq datasets were aligned and quantified using the CellRanger software package (version 4.0.0) and subjected to Seurat 3(version 3.2.3) (Hao et al., 2021) for further analysis. To pass quality control, cells were required to have less than 60000 raw reads mapping to nuclear genes, at least 2000 genes detected, less than 10% of mapped reads mapping to mitochondrial genes. Finally, 7977 cells passed the quality control. To rule out the effects of cell cycle variances, we performed a simple linear regression against the cell cycle score calculated by CellCycleScoring. Then, PCA was performed by RunPCA using 2000 highly variable genes and the top 20 PCs were used for UMAP analysis. Clusters were detected using FindClusters with setting parameters dims = 1:20, resolution = 0.08. Up-regulation genes were identified for each cluster using Wilcoxon rank sum test with parameter ‘min.pct = 0.5, logfc.threshold = 0.25’ implemented in Seurat. And the Gene ontology-enrichment analysis (biological processes) were performed for the up-regulation genes of each cluster by clusterProfiler with BH adjusted p-value cutoff =0.05 (version 3.14.3) (Yu et al., 2012a).

### QUANTIFICATION AND STATISTICAL ANALYSIS

Data analysis were performed using GraphPad Prism. All data are expressed as mean and exact n numbers for each dataset are detailed in the figure legends. All statistical analyses were performed by independent-sample student t test and Mann-Whitney U test (SPSS software). NS, not significant; *P< 0.05; **P< 0.01; ***P< 0.001.

## REFERENCES

Allman, D., and Pillai, S. (2008). Peripheral B cell subsets. Curr Opin Immunol 20, 149–157.

Aspland, S.E., Bendall, H.H., and Murre, C. (2001). The role of E2A-PBX1 in leukemogenesis. Oncogene 20, 5708–5717.

Bain, G., Maandag, E.C., Izon, D.J., Amsen, D., Kruisbeek, A.M., Weintraub, B.C., Krop, I., Schlissel, M.S., Feeney, A.J., van Roon, M., and et al. (1994). E2A proteins are required for proper B cell development and initiation of immunoglobulin gene rearrangements. Cell 79, 885–892.

Barber, C.L., Montecino-Rodriguez, E., and Dorshkind, K. (2011). Reduced production of B-1-specified common lymphoid progenitors results in diminished potential of adult marrow to generate B-1 cells. Proc Natl Acad Sci U S A 108, 13700–13704.

Bassing, C.H., Swat, W., and Alt, F.W. (2002). The mechanism and regulation of chromosomal V(D)J recombination. Cell 109 Suppl, S45–55.

Baumgarth, N. (2011). The double life of a B-1 cell: self-reactivity selects for protective effector functions. Nat Rev Immunol 11, 34–46.

Bolotin, D.A., Poslavsky, S., Mitrophanov, I., Shugay, M., Mamedov, I.Z., Putintseva, E.V., and Chudakov, D.M. (2015). MiXCR: software for comprehensive adaptive immunity profiling. Nat Methods 12, 380–381.

Burrows, P.D., Stephan, R.P., Wang, Y.H., Lassoued, K., Zhang, Z., and Cooper, M.D. (2002). The transient expression of pre-B cell receptors governs B cell development. Semin Immunol 14, 343–349.

Cerutti, A., Cols, M., and Puga, I. (2013). Marginal zone B cells: virtues of innate-like antibody-producing lymphocytes. Nat Rev Immunol 13, 118–132.

Cho, S.K., Webber, T.D., Carlyle, J.R., Nakano, T., Lewis, S.M., and Zuniga-Pflucker, J.C. (1999). Functional characterization of B lymphocytes generated in vitro from embryonic stem cells. Proc Natl Acad Sci U S A 96, 9797–9802.

Dai, X., Chen, Y., Di, L., Podd, A., Li, G., Bunting, K.D., Hennighausen, L., Wen, R., and Wang, D. (2007). Stat5 is essential for early B cell development but not for B cell maturation and function. J Immunol 179, 1068–1079.

De Silva, N.S., and Klein, U. (2015). Dynamics of B cells in germinal centres. Nat Rev Immunol 15, 137–148.

Dengler, H.S., Baracho, G.V., Omori, S.A., Bruckner, S., Arden, K.C., Castrillon, D.H., DePinho, R.A., and Rickert, R.C. (2008). Distinct functions for the transcription factor Foxo1 at various stages of B cell differentiation. Nat Immunol 9, 1388–1398.

French, A., Yang, C.T., Taylor, S., Watt, S.M., and Carpenter, L. (2015). Human induced pluripotent stem cell-derived B lymphocytes express sIgM and can be generated via a hemogenic endothelium intermediate. Stem Cells Dev 24, 1082–1095.

Ghosn, E.E., Yamamoto, R., Hamanaka, S., Yang, Y., Herzenberg, L.A., Nakauchi, H., and Herzenberg, L.A. (2012). Distinct B-cell lineage commitment distinguishes adult bone marrow hematopoietic stem cells. Proc Natl Acad Sci U S A 109, 5394–5398.

Gong, S., and Nussenzweig, M.C. (1996). Regulation of an early developmental checkpoint in the B cell pathway by Ig beta. Science 272, 411–414.

Grawunder, U., Leu, T.M., Schatz, D.G., Werner, A., Rolink, A.G., Melchers, F., and Winkler, T.H. (1995). Down-regulation of RAG1 and RAG2 gene expression in preB cells after functional immunoglobulin heavy chain rearrangement. Immunity 3, 601–608.

Guo, R., Hu, F., Weng, Q., Lv, C., Wu, H., Liu, L., Li, Z., Zeng, Y., Bai, Z., Zhang, M., et al. (2020). Guiding T lymphopoiesis from pluripotent stem cells by defined transcription factors. Cell Res 30, 21–33.

Hadland, B.K., Varnum-Finney, B., Mandal, P.K., Rossi, D.J., Poulos, M.G., Butler, J.M., Rafii, S., Yoder, M.C., Yoshimoto, M., and Bernstein, I.D. (2017). A Common Origin for B-1a and B-2 Lymphocytes in Clonal Pre-Hematopoietic Stem Cells. Stem Cell Reports 8, 1563–1572.

Hao, Y., Hao, S., Andersen-Nissen, E., Mauck, W.M., 3rd, Zheng, S., Butler, A., Lee, M.J., Wilk, A.J., Darby, C., Zager, M., et al. (2021). Integrated analysis of multimodal single-cell data. Cell 184, 3573–3587 e3529.

Hardy, R.R., Carmack, C.E., Shinton, S.A., Kemp, J.D., and Hayakawa, K. (1991). Resolution and characterization of pro-B and pre-pro-B cell stages in normal mouse bone marrow. J Exp Med 173, 1213–1225.

Hirose, Y., Kiyoi, H., Itoh, K., Kato, K., Saito, H., and Naoe, T. (2001). B-cell precursors differentiated from cord blood CD34+ cells are more immature than those derived from granulocyte colony-stimulating factor-mobilized peripheral blood CD34+ cells. Immunology 104, 410–417.

Hu, Y., Zhang, Z., Kashiwagi, M., Yoshida, T., Joshi, I., Jena, N., Somasundaram, R., Emmanuel, A.O., Sigvardsson, M., Fitamant, J., et al. (2016). Superenhancer reprogramming drives a B-cell-epithelial transition and high-risk leukemia. Genes Dev 30, 1971–1990.

Kerner, J.D., Appleby, M.W., Mohr, R.N., Chien, S., Rawlings, D.J., Maliszewski, C.R., Witte, O.N., and Perlmutter, R.M. (1995). Impaired expansion of mouse B cell progenitors lacking Btk. Immunity 3, 301–312.

Kitajima, K., Kawaguchi, M., Miyashita, K., Nakajima, M., Kanokoda, M., and Hara, T. (2015). Efficient production of T cells from mouse pluripotent stem cells by controlled expression of Lhx2. Genes Cells 20, 720–738.

Kitamura, D., Kudo, A., Schaal, S., Müller, W., Melchers, F., and Rajewsky, K. (1992). A critical role of λ5 protein in B cell development. Cell 69, 823–831.

Kitamura, D., Roes, J., Kuhn, R., and Rajewsky, K. (1991). A B cell-deficient mouse by targeted disruption of the membrane exon of the immunoglobulin mu chain gene. Nature 350, 423–426.

Kobayashi, M., Shelley, W.C., Seo, W., Vemula, S., Lin, Y., Liu, Y., Kapur, R., Taniuchi, I., and Yoshimoto, M. (2014). Functional B-1 progenitor cells are present in the hematopoietic stem cell-deficient embryo and depend on Cbfbeta for their development. Proc Natl Acad Sci U S A 111, 12151–12156.

Kobayashi, M., Tarnawsky, S.P., Wei, H., Mishra, A., Azevedo Portilho, N., Wenzel, P., Davis, B., Wu, J., Hadland, B., and Yoshimoto, M. (2019). Hemogenic Endothelial Cells Can Transition to Hematopoietic Stem Cells through a B-1 Lymphocyte-Biased State during Maturation in the Mouse Embryo. Stem Cell Reports 13, 21–30.

Li, Y.S., Wasserman, R., Hayakawa, K., and Hardy, R.R. (1996). Identification of the earliest B lineage stage in mouse bone marrow. Immunity 5, 527–535.

Lin, H., and Grosschedl, R. (1995). Failure of B-cell differentiation in mice lacking the transcription factor EBF. Nature 376, 263–267.

Lin, Y., Kobayashi, M., Azevedo Portilho, N., Mishra, A., Gao, H., Liu, Y., Wenzel, P., Davis, B., Yoder, M.C., and Yoshimoto, M. (2019). Long-Term Engraftment of ESC-Derived B-1 Progenitor Cells Supports HSC-Independent Lymphopoiesis. Stem Cell Reports 12, 572–583.

Liu, G.J., Cimmino, L., Jude, J.G., Hu, Y., Witkowski, M.T., McKenzie, M.D., Kartal-Kaess, M., Best, S.A., Tuohey, L., Liao, Y., et al. (2014). Pax5 loss imposes a reversible differentiation block in B-progenitor acute lymphoblastic leukemia. Genes Dev 28, 1337–1350.

Lu, Y.F., Cahan, P., Ross, S., Sahalie, J., Sousa, P.M., Hadland, B.K., Cai, W., Serrao, E., Engelman, A.N., Bernstein, I.D., and Daley, G.Q. (2016). Engineered Murine HSCs Reconstitute Multi-lineage Hematopoiesis and Adaptive Immunity. Cell Rep 17, 3178–3192.

Ma, S., Pathak, S., Mandal, M., Trinh, L., Clark, M.R., and Lu, R. (2010). Ikaros and Aiolos inhibit pre-B-cell proliferation by directly suppressing c-Myc expression. Mol Cell Biol 30, 4149–4158.

Ma, S., Pathak, S., Trinh, L., and Lu, R. (2008). Interferon regulatory factors 4 and 8 induce the expression of Ikaros and Aiolos to down-regulate pre-B-cell receptor and promote cell-cycle withdrawal in pre-B-cell development. Blood 111, 1396–1403.

Mackay, F., and Schneider, P. (2009). Cracking the BAFF code. Nat Rev Immunol 9, 491–502.

Martensson, A., Argon, Y., Melchers, F., Dul, J.L., and Martensson, I.L. (1999). Partial block in B lymphocyte development at the transition into the pre-B cell receptor stage in Vpre-B1-deficient mice. Int Immunol 11, 453–460.

Martin, F., Oliver, A.M., and Kearney, J.F. (2001). Marginal zone and B1 B cells unite in the early response against T-independent blood-borne particulate antigens. Immunity 14, 617–629.

McKearn, J.P., Baum, C., and Davie, J.M. (1984). Cell surface antigens expressed by subsets of pre-B cells and B cells. J Immunol 132, 332–339.

Melchers, F. (2015). Checkpoints that control B cell development. J Clin Invest 125, 2203–2210.

Muljo, S.A., and Schlissel, M.S. (2003). A small molecule Abl kinase inhibitor induces differentiation of Abelson virus-transformed pre-B cell lines. Nat Immunol 4, 31–37.

Ng, A.P., Coughlan, H.D., Hediyeh-Zadeh, S., Behrens, K., Johanson, T.M., Low, M.S.Y., Bell, C.C., Gilan, O., Chan, Y.C., Kueh, A.J., et al. (2020). An Erg-driven transcriptional program controls B cell lymphopoiesis. Nat Commun 11, 3013.

Ohkawara, J.I., Ikebuchi, K., Fujihara, M., Sato, N., Hirayama, F., Yamaguchi, M., Mori, K.J., and Sekiguchi, S. (1998). Culture system for extensive production of CD19+IgM+ cells by human cord blood CD34+ progenitors. Leukemia 12, 764–771.

Okuno, Y., and Yuki, H. (2012). PU.1 is a tumor suppressor for B cell malignancies. Oncotarget 3, 1495–1496.

Opstelten, D., and Osmond, D.G. (1983). Pre-B cells in mouse bone marrow: immunofluorescence stathmokinetic studies of the proliferation of cytoplasmic mu-chain-bearing cells in normal mice. J Immunol 131, 2635–2640.

Osmond, D.G., and Owen, J.J. (1984). Pre-B cells in bone marrow: size distribution profile, proliferative capacity and peanut agglutinin binding of cytoplasmic mu chain-bearing cell populations in normal and regenerating bone marrow. Immunology 51, 333–342.

Osmond, D.G., Rolink, A., and Melchers, F. (1998). Murine B lymphopoiesis: towards a unified model. Immunol Today 19, 65–68.

Pappu, R., Cheng, A.M., Li, B., Gong, Q., Chiu, C., Griffin, N., White, M., Sleckman, B.P., and Chan, A.C. (1999). Requirement for B cell linker protein (BLNK) in B cell development. Science 286, 1949–1954.

Park, Y.H., and Osmond, D.G. (1987). Phenotype and proliferation of early B lymphocyte precursor cells in mouse bone marrow. J Exp Med 165, 444–458.

Pearson, S., Cuvertino, S., Fleury, M., Lacaud, G., and Kouskoff, V. (2015). In vivo repopulating activity emerges at the onset of hematopoietic specification during embryonic stem cell differentiation. Stem Cell Reports 4, 431–444.

Pereira, C.F., Chang, B., Qiu, J., Niu, X., Papatsenko, D., Hendry, C.E., Clark, N.R., Nomura-Kitabayashi, A., Kovacic, J.C., Ma’ayan, A., et al. (2013). Induction of a hemogenic program in mouse fibroblasts. Cell Stem Cell 13, 205–218.

Potocnik, A.J., Nerz, G., Kohler, H., and Eichmann, K. (1997). Reconstitution of B cell subsets in Rag deficient mice by transplantation of in vitro differentiated embryonic stem cells. Immunol Lett 57, 131–137.

Reynaud, D., Demarco, I.A., Reddy, K.L., Schjerven, H., Bertolino, E., Chen, Z., Smale, S.T., Winandy, S., and Singh, H. (2008). Regulation of B cell fate commitment and immunoglobulin heavy-chain gene rearrangements by Ikaros. Nat Immunol 9, 927–936.

Richter, K., Pinto do, O.P., Hagglund, A.C., Wahlin, A., and Carlsson, L. (2003). Lhx2 expression in hematopoietic progenitor/stem cells in vivo causes a chronic myeloproliferative disorder and altered globin expression. Haematologica 88, 1336–1347.

Rolink, A., Grawunder, U., Winkler, T.H., Karasuyama, H., and Melchers, F. (1994). IL-2 receptor alpha chain (CD25, TAC) expression defines a crucial stage in pre-B cell development. Int Immunol 6, 1257–1264.

Scheuermann, R.H., and Racila, E. (1995). CD19 antigen in leukemia and lymphoma diagnosis and immunotherapy. Leuk Lymphoma 18, 385–397.

Scott, E.W., Simon, M.C., Anastasi, J., and Singh, H. (1994). Requirement of transcription factor PU.1 in the development of multiple hematopoietic lineages. Science 265, 1573–1577.

Shugay, M., Bagaev, D.V., Turchaninova, M.A., Bolotin, D.A., Britanova, O.V., Putintseva, E.V., Pogorelyy, M.V., Nazarov, V.I., Zvyagin, I.V., Kirgizova, V.I., et al. (2015). VDJtools: Unifying Post-analysis of T Cell Receptor Repertoires. PLoS Comput Biol 11, e1004503.

Smith, F.L., and Baumgarth, N. (2019). B-1 cell responses to infections. Curr Opin Immunol 57, 23–31.

Stebegg, M., Kumar, S.D., Silva-Cayetano, A., Fonseca, V.R., Linterman, M.A., and Graca, L. (2018). Regulation of the Germinal Center Response. Front Immunol 9, 2469.

Stik, G., and Graf, T. (2018). Hoxb5, a Trojan horse to generate T cells. Nat Immunol 19, 210–212.

Sugimura, R., Jha, D.K., Han, A., Soria-Valles, C., da Rocha, E.L., Lu, Y.F., Goettel, J.A., Serrao, E., Rowe, R.G., Malleshaiah, M., et al. (2017). Haematopoietic stem and progenitor cells from human pluripotent stem cells. Nature 545, 432–438.

Swaminathan, S., Huang, C., Geng, H., Chen, Z., Harvey, R., Kang, H., Ng, C., Titz, B., Hurtz, C., Sadiyah, M.F., et al. (2013). BACH2 mediates negative selection and p53-dependent tumor suppression at the pre-B cell receptor checkpoint. Nat Med 19, 1014–1022.

Thompson, E.C., Cobb, B.S., Sabbattini, P., Meixlsperger, S., Parelho, V., Liberg, D., Taylor, B., Dillon, N., Georgopoulos, K., Jumaa, H., et al. (2007). Ikaros DNA-binding proteins as integral components of B cell developmental-stage-specific regulatory circuits. Immunity 26, 335–344.

Tonegawa, S. (1983). Somatic generation of antibody diversity. Nature 302, 575–581.

Urbanek, P., Wang, Z.Q., Fetka, I., Wagner, E.F., and Busslinger, M. (1994). Complete block of early B cell differentiation and altered patterning of the posterior midbrain in mice lacking Pax5/BSAP. Cell 79, 901–912.

Vale, A.M., and Schroeder, H.W., Jr. (2010). Clinical consequences of defects in B-cell development. J Allergy Clin Immunol 125, 778–787.

Villeval, J.L., Testa, U., Vinci, G., Tonthat, H., Bettaieb, A., Titeux, M., Cramer, P., Edelman, L., Rochant, H., Bretongorius, J., and Vainchenker, W. (1985). Carbonic Anhydrase-I Is an Early Specific Marker of Normal Human Erythroid-Differentiation. Blood 66, 1162–1170.

Vinuesa, C.G., Linterman, M.A., Goodnow, C.C., and Randall, K.L. (2010). T cells and follicular dendritic cells in germinal center B-cell formation and selection. Immunol Rev 237, 72–89.

Wang, T., Lv, C., Hu, F., Liu, L., and Wang, J. (2020). Two-step protocol for regeneration of immunocompetent T cells from mouse pluripotent stem cells. Blood Science 2, 79–88.

Xu, Y., Baldassare, M., Fisher, P., Rathbun, G., Oltz, E.M., Yancopoulos, G.D., Jessell, T.M., and Alt, F.W. (1993). LH-2: a LIM/homeodomain gene expressed in developing lymphocytes and neural cells. Proc Natl Acad Sci U S A 90, 227–231.

Yoshimoto, M., Montecino-Rodriguez, E., Ferkowicz, M.J., Porayette, P., Shelley, W.C., Conway, S.J., Dorshkind, K., and Yoder, M.C. (2011). Embryonic day 9 yolk sac and intra-embryonic hemogenic endothelium independently generate a B-1 and marginal zone progenitor lacking B-2 potential. Proc Natl Acad Sci U S A 108, 1468–1473.

Yu, G., Wang, L.G., Han, Y., and He, Q.Y. (2012a). clusterProfiler: an R package for comparing biological themes among gene clusters. OMICS 16, 284–287.

Yu, Y., Wang, J., Khaled, W., Burke, S., Li, P., Chen, X., Yang, W., Jenkins, N.A., Copeland, N.G., Zhang, S., and Liu, P. (2012b). Bcl11a is essential for lymphoid development and negatively regulates p53. J Exp Med 209, 2467–2483.

Zheng, Y., Yu, M., Podd, A., Yuan, L., Newman, D.K., Wen, R., Arepally, G., and Wang, D. (2013). Critical role for mouse marginal zone B cells in PF4/heparin antibody production. Blood 121, 3484–3492.

Zhou, F., Li, X., Wang, W., Zhu, P., Zhou, J., He, W., Ding, M., Xiong, F., Zheng, X., Li, Z., et al. (2016). Tracing haematopoietic stem cell formation at single-cell resolution. Nature 533, 487–492.

Zouali, M., and Richard, Y. (2011). Marginal zone B-cells, a gatekeeper of innate immunity. Front Immunol 2, 63.

