## Supplemental Figures for "Regeneration of humoral immunity from pluripotent stem cells by defined transcription factors"

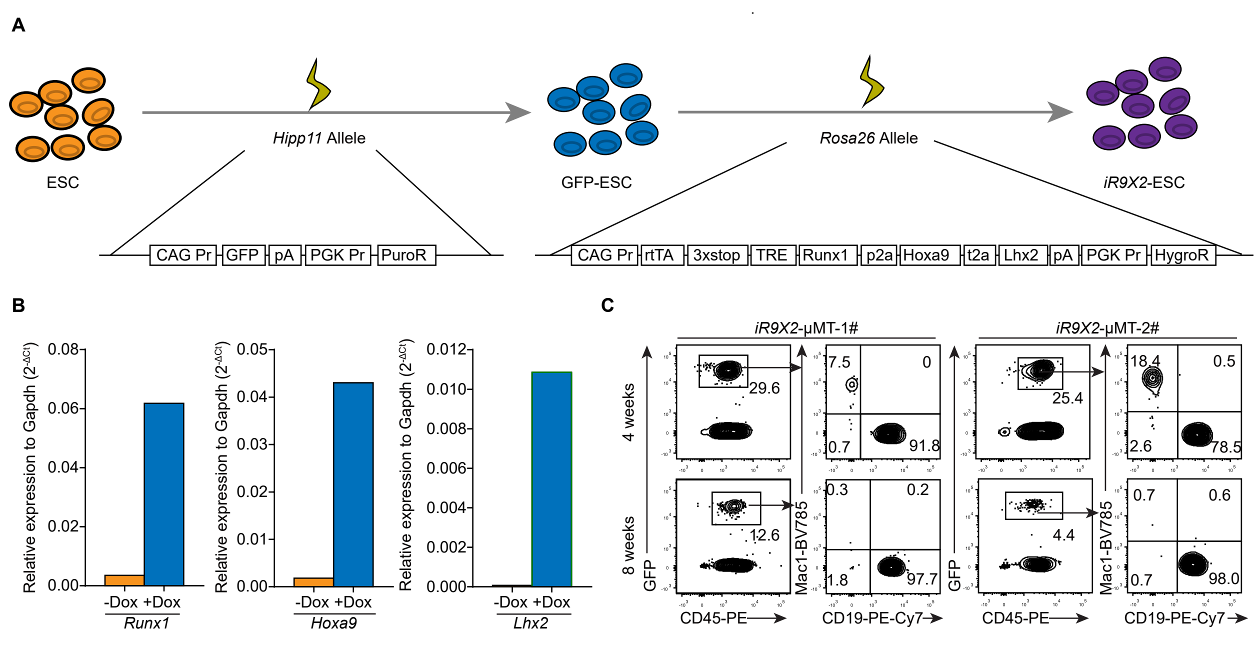


**Figure S1. Reconstitution of functional B cells *in vivo* from *iR9X2*-ESC, Related to Figure 1.** (A) Schematic diagram of establishing the *iR9X2*-ESC cell line. A CAG Pr-GFP-PGK Pr-PuroR cassette was inserted into the *Hipp11* locus of mouse ESC by homologous recombination to form GFP-ESC clones which were selected by puromycine (1 μg/ mL). Next, a CAG Pr-rtTA-3 × Stop-TRE-Runx1-p2a-Hoxa9-t2a-Lhx2-pA-PGK Pr-HygroR cassette was inserted into the *Rosa26* locus of GFP-ESC to form *iR9X2*-ESC clones which were selected by hygromycin B (150 μg/ml). (B) Q-PCR analysis of the expression of *Runx1*, *Hoxa9,* and *Lhx2* in *iR9X2*-ESC after doxycycline induction. (C) *iR9X2*-ESCs-derived $GFP^+CD45^+Mac1^+$ myeloid cells and $GFP^+CD45^+CD19^+$ B cells in the PB of μMT mice were analyzed by flow cytometry 4 weeks and 8 weeks after transplantation. Five million iHECs-derived hematopoietic progenitors were transplanted into each sublethally irradiated μMT mouse. The mice were fed with water containing doxycycline (1 mg/mL) to induce the generation of B lymphocytes. Two representative mice from five independent experiments are shown.

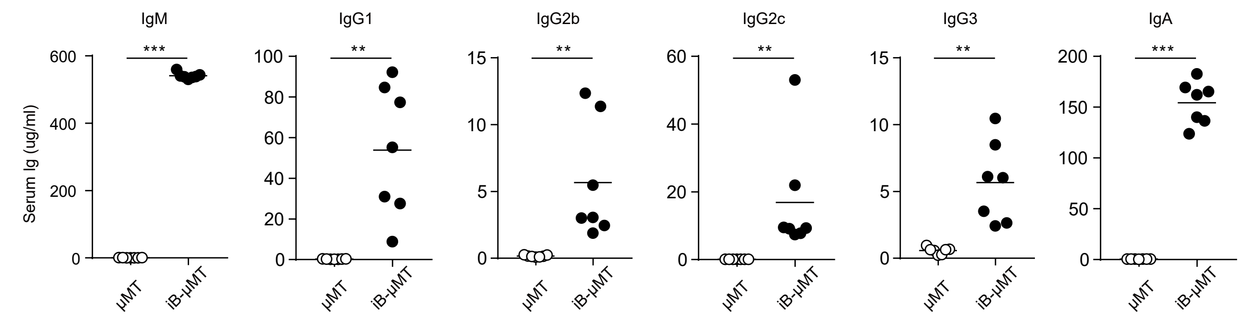


**Figure S2. Long term serum antibodies levels in iB-μMT mice, Related to Figure 2.** Sera were collected from iB-μMT mice (18 to 40 weeks after transplantation) and μMT mice (n=7 per group). The different isotypes of antibodies (IgM /IgG1 /IgG2b /IgG2c /IgG3 /IgA) were measured by ELISA. Each symbol represents an individual mouse; small horizontal lines indicate the mean. $^*P < 0.05$, $^{**}P < 0.01$, and $^{***}P < 0.001$ (Independent-sample student t test and Mann-Whitney U test).

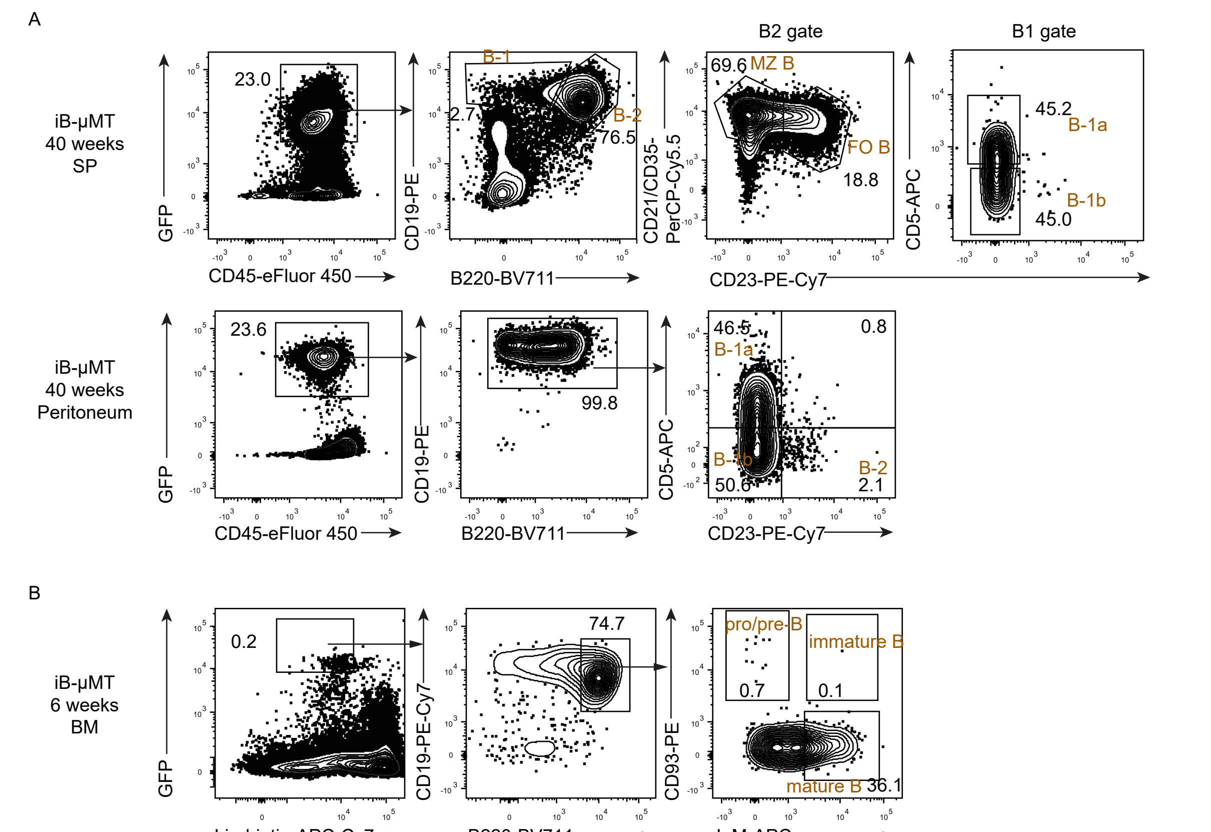


**Figure S3. The regenerative B cell subsets in iB-μMT mice at week 6 and week 40 after transplantation, Related to Figure 3.** (A) All major mature B cell subsets were analyzed by flow cytometry at week 40 after transplantation. Immuno-phenotypic plots of B-1a, B-1b, FO B, and MZ B cells in the spleen of iB-μMT mice were depicted. And immune-phenotypic plots of B-1a, B-1b, and B-2 cells in the peritoneal cavity of iB-μMT mice were depcited. Data from one representative mouse are shown. (B) Bone marrow cells were isolated from iB-μMT mice 6 weeks after transplantation, and flow cytometry plots of pro/pre-B cells, immature B cells, and mature B cells were depicted. Data from one representative mouse are shown. $Lin^-$ was defined as $Ter119^-$ $Mac1^-$ $Gr1^-$ $NK1.1^-$ $CD3^-$ $CD4^-$ $CD8^-$.

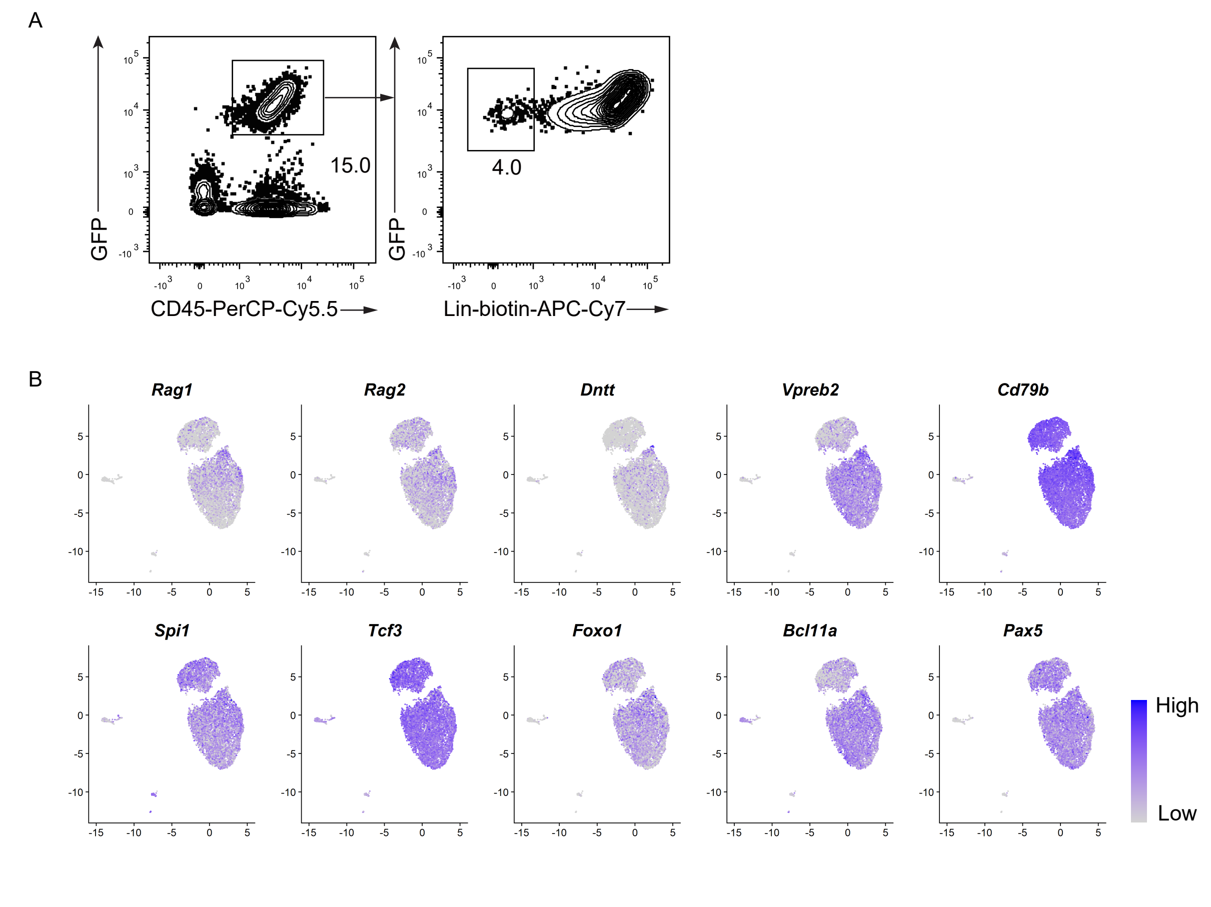


**Figure S4. Sorting gates of bone marrow regenerative progenitors and UMAP analysis of the expression patterns of selected early B progenitors related genes, Related to Figure 4.** (A) Bone marrow cells were sorted based on $GFP^+CD45^+Lin^-$ surface markers for single cell sequencing at day7.5 after transplantation, $Lin^-$ was defined as $CD3^-$ $CD4^-$ $CD8^-$ $Ter119^-$ $Mac1^-$ $Gr1^-$ $NK1.1^-$. (B) Single-cell RNA-seq analysis (7977 cells) showing UMAP plots of VDJ recombination related enzymes genes (*Rag1*, *Rag2,* and *Dntt*), pre-BCR complex related genes (*Vpreb2* and *Cd79b*), and early B cell development related transcription factors (*Spi1*, *Tcf3*, *Foxo1*, *Bcl11a*, and *Pax5* ). The expression value of each gene was converted by log2 and illustrated by FeaturePlots of Seurat3.

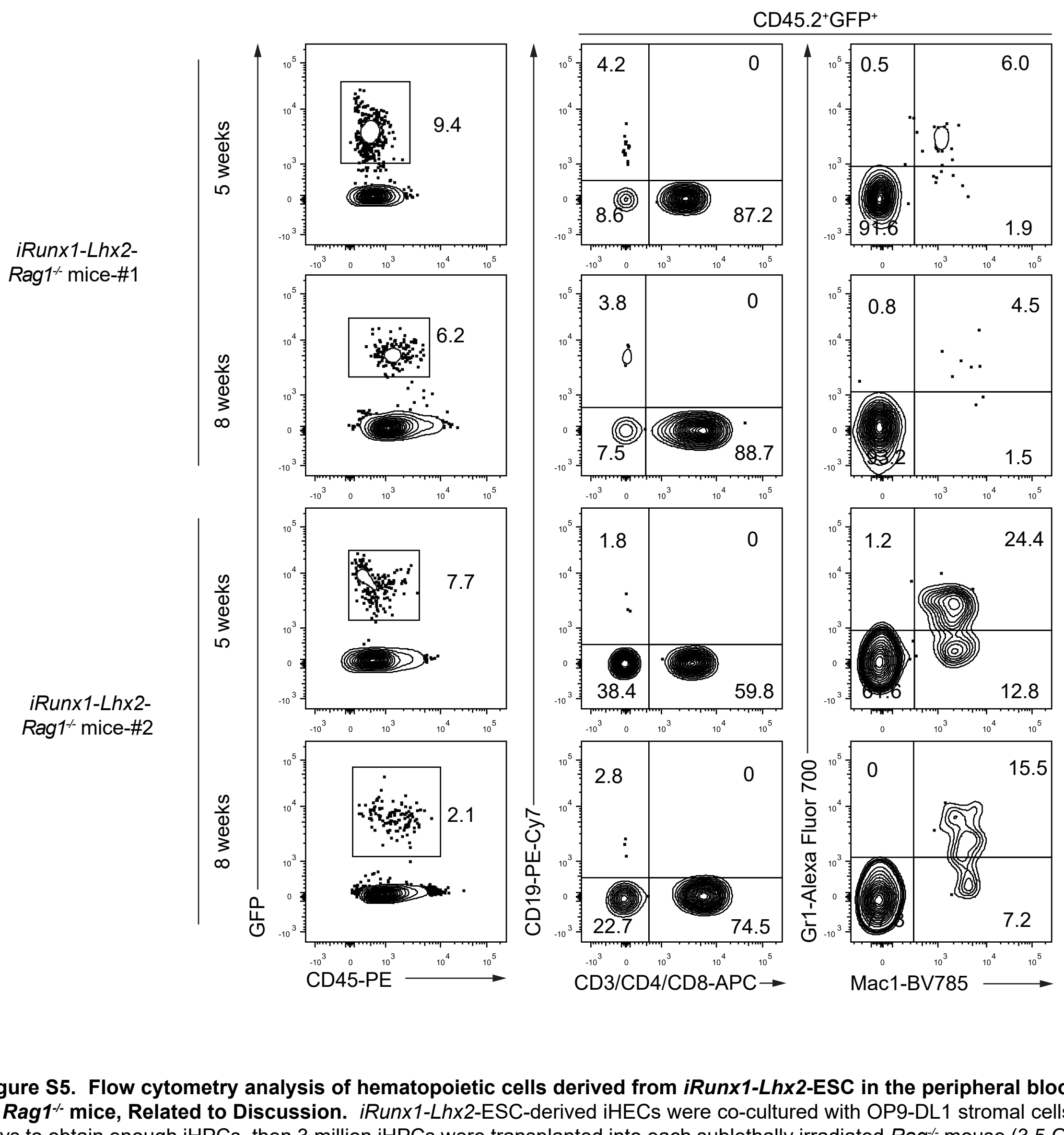


**Figure S5. Flow cytometry analysis of hematopoietic cells derived from *iRunx1-Lhx2*-ESC in the peripheral blood (PB) of *Rag1*$^{-/-}$ mice, Related to Discussion.** *iRunx1-Lhx2*-ESC-derived iHECs were co-cultured with OP9-DL1 stromal cells for 15 days to obtain enough iHPCs, then 3 million iHPCs were transplanted into each sublethally irradiated *Rag*$^{-/-}$ mouse (3.5 Gy). The mice were fed with water containing doxycycline (1 mg/mL). Hematopoietic cells ($Mac1^+$ myeloid cells, $CD3/CD4/CD8^+$ T cells, and $CD19^+$ B cells) in the peripheral blood of *Rag*$^{-/-}$ recipients were analyzed by flow cytometry at week 5 and week 8 after transplantation. Data from two representative mice are shown.
